# Calorie Restriction in combination with GABA: improved glycaemic control in type 2 diabetes mouse model

**DOI:** 10.1101/2021.05.12.443226

**Authors:** Nirali Rathwa, Nishant Parmar, Sayantani Pramanik Palit, Roma Patel, Ravi Shankar Bhaskaran, A. V. Ramachandran, Rasheedunnisa Begum

## Abstract

**Aim:** Dysfunctional adipocytes and β-cells are crucial in the advancement of obesity-induced type 2 diabetes (T2D). Calorie restriction (CR) improves insulin sensitivity and fasting blood glucose (FBG) levels, while γ-aminobutyric acid (GABA), a neurotransmitter secreted by β-cells, exerts regenerative effects.

**Methods:** High-fat diet (HFD) and streptozotocin (STZ) induced T2D mouse model was used to assess the effect of CR and GABA combination therapy. The mice were fed with CR diet (30% reduction of HFD) and treated with GABA (2.5 mg/kg i.p) daily for 5 weeks. Assessment of FBG, body weight (BW), insulin sensitivity, glucose tolerance, and estimation of plasma insulin and lipid profile were carried out by standard methods. The mRNA expression of target genes [liver (glucoregulatory enzymes), adipose tissue (lipid metabolism markers) and skeletal muscle (mitochondrial biogenesis markers)] was assessed by SYBR Green-qPCR. Oxygen consumption rate for mitochondrial complex I, II and III was measured in skeletal muscle using Oxytherm Clark-type oxygen electrode. The effect of monotherapies and combination treatment on pancreatic β-cell regeneration and apoptosis was analysed by IHC.

**Results:** CR+GABA group showed significant reduction in FBG levels, improved lipid profile and whole-body insulin responsiveness as indicated by increased insulin levels, insulin sensitivity and glucose tolerance compared to the monotherapies. The transcript expression profile of glucoregulatory enzymes in the liver showed significant reduction in *G6Pase, PEPCK, Glycogen Phosphorylase* and increased *GCK* expression in CR+GABA group as compared to the monotherapies The *ACC and ATG expression was* up-regulated in adipose tissue; and significant increase in *SIRT-1, PGC-1*α and *TFAM* expression with up-regulated mitochondrial complex I and III activities was observed in CR+GABA group as compared to the monotherapies. Percentage of BrdU/Insulin and PDX1/Ngn3/Insulin co-positive cells was significantly higher in CR+GABA treated group than the monotherapy and HFD+STZ groups. Finally, there was a significant reduction in TUNEL/Insulin co-positive cells suggesting reduced apoptosis in CR+GABA treated group.

**Conclusion:** Our results suggest that CR in combination with GABA ameliorates T2D in HFD+STZ treated mice, by GABA induced β-cell regeneration and CR mediated insulin sensitivity.

## 1. Introduction

Dysfunctional adipocytes and β-cells are involved in advancing obesity-induced type 2 diabetes (T2D) [1]. Several aspects regulate the progression of T2D, such as genetic predisposition [2-9], excess calorie intake [1, 10] and sedentary lifestyle [10], all of which cause persistent hyperglycemia and eventual β-cell apoptosis. Even though there have been many recent management therapies for T2D, none can withstand the worsening condition of β-cell loss. This wide gap has triggered the search for alternative remedies that can regenerate β-cells from either stem cells or existing β-cells and bring about insulin sensitivity.

Calorie restriction (CR), an effective dietary intervention for T2D, has been associated with longevity. CR reduces caloric intake, typically by 20–40% of ad libitum consumption, while maintaining adequate intake of protein and micronutrients to avoid malnutrition [13]. CR attenuates the degree of oxidative stress [14, 15] and increases the transcript levels of genes involved in mitochondrial function and biogenesis [15]. It also improves insulin sensitivity [13, 14], FBG, other cardiometabolic risk factors, and reduces pro-inflammatory adipokines and total cholesterol [16].

γ-aminobutyric acid (GABA) has emerged as a new antidiabetic dietary supplement. GABA, a neurotransmitter, is secreted by the central nervous system (CNS) and pancreatic β-cells. It mediates protective and regenerative effects by reducing β-cell apoptosis and increasing its replication rate [17, 18]. GABA exerts its action via two primary receptors GABAA and GABAB, which regulates insulin and glucagon secretion in the pancreatic islets [19]. GABA has shown its type 1 diabetes ameliorating effect by suppressing Treg cells in long-term therapy where GABA has demonstrated induction of β-cell regeneration by proliferation and transdifferentiation [20, 21].

We have investigated the combinatorial effect of CR and GABA in the T2D mouse model induced by high-fat diet (HFD) + streptozotocin (HFD+STZ). To evaluate the effect of CR and GABA on multiple targets and regulatory pathways, we assessed i) insulin sensitivity, glucose tolerance, and metabolic profile, ii) transcript levels of key genes involved in hepatic glucoregulation and lipid metabolism in adipose tissue (AT) and mitochondrial biogenesis in skeletal muscle (SK), iii) ETC complex I, II and III activities in SK, and iv) regenerative markers in β-cells.

## 2. Materials and Methods

### 2.1 Animals and experimental strategy

#### 2.1.1 Ethics statement

The experimental procedures were approved by the Institutional Ethical Committee for Animal Research (IECHR), Faculty of Science, The Maharaja Sayajirao University of Baroda, Vadodara, Gujarat, India (FS/IECHR/2016-9).

#### 2.1.2 Animals

Male C57BL/6 mice (∼8-10 weeks old, approximately 20-22 grams) were procured from ACTREC, Mumbai. They were housed at 23.0±1°C with a 12-hrs light/dark cycle and acclimatised for one week. These animals had free access to standard chow/HFD/CR diet (Keval Sales Corporation, Vadodara, India) and water.

#### 2.1.3 Development of T2D mouse model

The mice were divided into two groups. The control group (n=6) was fed with a normal chow diet (NCD). Obesity can induce multiple damaging effects on peripheral tissues. Working on similar lines, mice (n=25) were fed with HFD for 20 weeks to induce obesity and insulin resistance. After 20 weeks, these mice received three consecutive doses of 40 mg/kg BW STZ i.p. (MP Biomedicals, India) to induce β-cell loss. The HFD+STZ model is characterised by a significantly increased weight (≥30 grams), hyperglycemia (FBG ≥300 mg/dL), hyperlipidemia, insulin resistance and β-cell loss [24].

#### 2.1.4 Treatment

The HFD+STZ treated animals were divided randomly into four groups (4-5 mice/group): 1. Diabetic control (HFD+STZ) 2. CR diet-fed (30% reduction of HFD) [22, 23] 3. GABA treated (2.5 mg/kg bw i.p) (Sigma–Aldrich, United States) [21] and 4. CR+GABA treated (30% reduction of HFD + GABA, 2.5 mg/kg bw i.p). Treatment was given daily for five weeks along with 100 mg/kg bw i.p BrdU (MP Biomedicals, India) on alternate days. CR diet comprised of 39% casein, 21% lard, 19 % sucrose, 16% vitamin-mineral premix, and 5% cellulose. The timeline is depicted in figure S1.

### 2.2 Metabolic and biochemical assessment

#### 2.2.1 Metabolic profiling

FBG was measured by tail snipping weekly using glucometer (TRUEresult, Nipro, India) along with body weight (BW) as well as food and water intake. At the end of the experiment, 1 ml of blood was collected from the orbital sinus for the biochemical assays after 5 hrs of fasting. Plasma was separated and used for assessing lipid profile [total cholesterol (TC), triglycerides (TG) and high-density lipoprotein (HDL)] by using commercial kits (Reckon Diagnostics P. Ltd, India). Friedewald’s (1972) formula was used for calculating low-density lipoprotein (LDL).

#### 2.2.2 Intraperitoneal Glucose Tolerance Test (IPGTT) and Intraperitoneal Insulin Sensitivity Test (IPIST)

IPGTT and IPIST were performed to evaluate glucose tolerance and insulin sensitivity at the end of 5 weeks of drug treatment. Mice were fasted for 6 hrs and injected with glucose (2g/kg BW i.p) or insulin (0.5U/kg BW i.p) for IPGTT and IPIST [25] respectively. Blood glucose levels were measured at the specified time points 0, 30, 60, 90 and 120 mins after glucose or insulin injection using glucometer.

#### 2.2.3 Assessment of insulin and C-peptide levels

The plasma levels of insulin and C-peptide in the experimental animals were measured using commercially available ELISA kits (RayBio, USA). All the plasma estimations were carried out in duplicate to ensure the coefficient of variation (CV) was below 10%.

### 2.3 Gene expression profiling

After sacrificing the mice, peripheral tissues such as liver, AT and SK were dissected and stored in RNAlater™ Stabilization Solution (Thermo Fisher Scientific, USA). Total RNA was extracted by the Trizol method as described previously [8]. The expression of targeted genes and *GAPDH* transcripts were monitored by LightCycler®480 Real-time PCR (Roche Diagnostics GmbH, Germany) using gene-specific primers (Eurofins, India) [Table S1]. Expression of *GAPDH* gene was used as a reference. Real-time PCR was performed as described previously [5].

### 2.4 Estimation of Oxygen Consumption Rate (OCR) of mitochondria

Skeletal muscle was isolated from left thigh of the mouse and mitochondria were isolated from using mitochondria isolation kit (Thermo Scientific TM, Catalog no. 89801) using manufacturer’s protocol. The isolated mitochondria were resuspended in mitochondria respiration buffer (80 mM KCl, 0.1% BSA, 50 mM HEPES, 2 mM MgCl2 and 2.5 mM KH2PO4; pH 7.2). Outer membrane integrity of the isolated mitochondria was evaluated by impermeability to exogenous cytochrome c which was constantly >95%. The activities of respiratory chain complexes I-III were recorded using 100 mM Pyruvate & 800 mM Malate (complex I), 1M Succinate (complex II) and 10 mM α-glycerophosphate (complex III) and, the protein concentration was estimated by Lowry *et al* method. OCR was determined by measuring the amount of oxygen (nmol) consumed, divided by the time elapsed (min) and the amount of protein present in the assay [26].

### 2.5 Pancreatic tissue preparation, Immunohistochemistry (IHC), assessment of β-cell regeneration and apoptosis

At the end of 5 weeks of drug treatment, animals were sacrificed, pancreatic tissues were fixed in 10% formalin and were processed for paraffin embedding. 5µm sections were cut from the paraffin-embedded blocks. To perform IHC, these sections were deparaffinised in100% xylene and washed serially in ethanol grades (100%, 95%, 80% and 70%) and proceeded with an antigen retrieval step (1N HCl at 37°C for 45 min). Subsequently, these sections were blocked with 5% donkey serum made in phosphate buffer saline (PBS) [Jackson ImmunoResearch Laboratories, Inc. USA], and antibodies were diluted in the blocking reagent. The details of the antibodies are as indicated in table S4. The sections were incubated with primary antibody for one hour at 37°C, washed with PBS, and incubated with secondary antibody for 45 min at 37°C. These sections were stained with anti-fade DAPI (Thermo Fisher Scientific, USA). The sections were covered with a coverslip. The sections were visualised under a confocal microscope (ZEISS LSM, Oberkochen, Germany) at 60X, and the images were processed by Image J (NIH, USA). Immunofluorescence staining (IFS) with anti-insulin and anti-BrdU was used to monitor β-cell proliferation. IFS with anti-insulin, anti-PAX-4 (paired box gene 4), and anti-ARX (Aristaless Related Homeobox) was carried out for α-cell to β-cell transdifferentiation while anti-insulin, anti-NGN-3 (Neurogenin 3) and anti-PDX-1 (pancreas/duodenum homeobox protein 1) were considered for β-cell neogenesis. IFS anti-insulin and TUNEL/anti-AIF (Apoptosis Inducing Factor) monitored β-cell death. Results obtained were expressed as the percentage of specified markers for regeneration and death.

### 2.6 Statistical analyses

The data was analysed and expressed as the mean ± SEM, and *p*<0.05 was considered statistically significant. The inter-group analysis was carried out by one-way ANOVA, followed by Tukey’s test for multiple group analysis. All the analyses were carried out using the Prism software 6 (GraphPad Prism, San Diego, CA, USA).

## 3. Results

### 3.1 HFD+STZ induced T2D mouse model establishment

After 20 weeks of HFD treatment, animals turned obese and insulin resistant. There was a significant increase in BW (Table S2a; Figures S1a&b). After two weeks of STZ administration, the mice turned hyperglycaemic as indicated by FBG levels ≥300 mg/dL (Table S 2b; Figures S1 c &d).

### 3.2 Metabolic and biochemical assessment

#### 3.2.1 Metabolic profiling

GABA treated group showed significantly reduced FBG levels (*p*<0.01), but no change was observed in BW, while CR monotherapy showed improvement in BW (*p*<0.05) but not in FBG levels. The BW (*p*<0.05) and FBG (*p*<0.01) levels of the CR+GABA treated group were significantly reduced at the end of 5 weeks of drug treatment compared to the HFD+STZ group, as shown in Figure 1.

**Figure 1.**
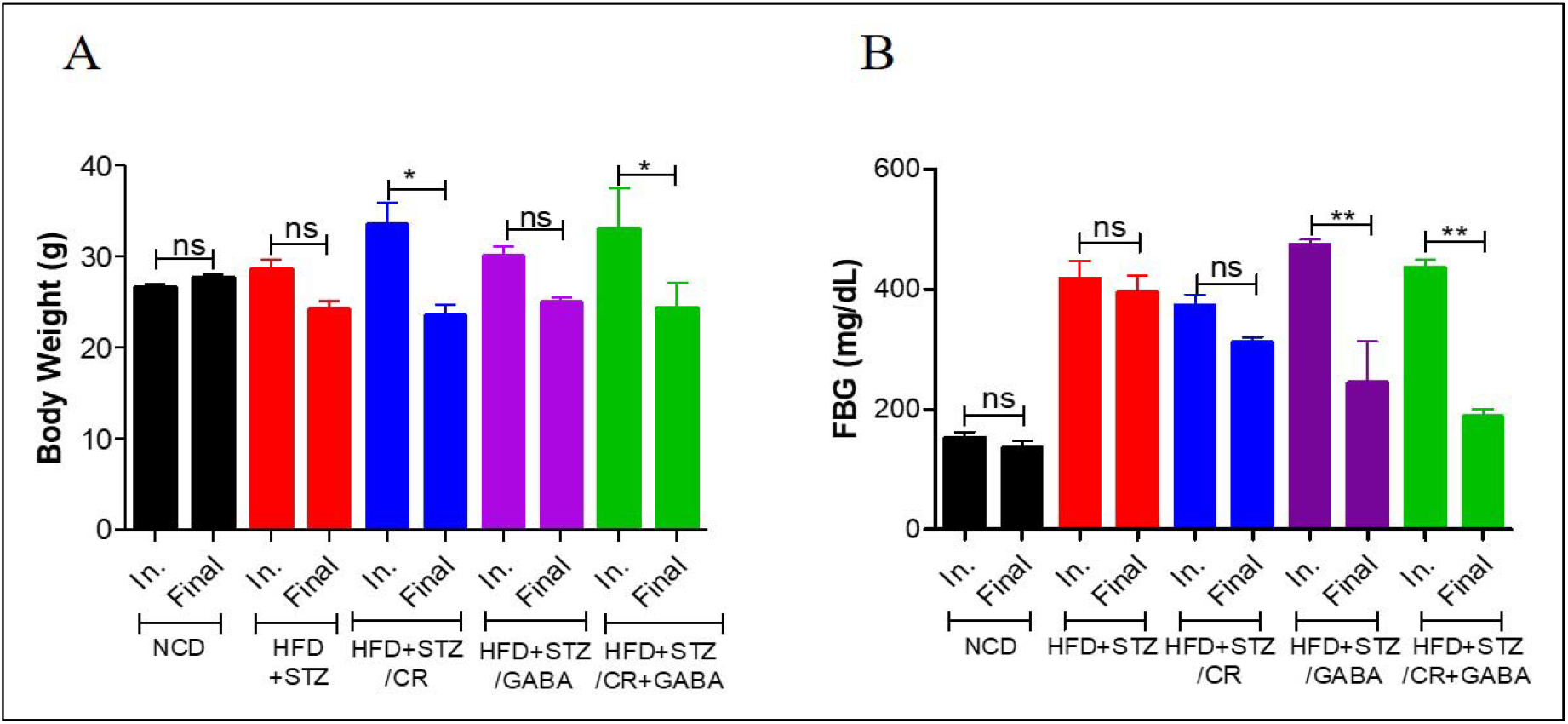
Body weight and fasting blood glucose levels. A) A significant decrease in BW was observed in the CR and CR+GABA treated groups compared to HFD+STZ. B) FBG levels in mice treated with GABA and CR+GABA were significantly reduced post-treatment (ns=non-significant, \**p*=0.05; \*\**p*<0.01, n=4-5/group).

#### 3.2.3 Lipid profiling

The GABA treated group showed no significant difference in the lipid profile (*p*>0.05), while CR fed mice (*p*<0.05) showed reduced TG levels as compared to HFD+STZ group. TC, TG and LDL levels were significantly reduced in CR+GABA treated group (*p*<0.05) as compared to HFD+STZ group (Figure 3).

#### 3.2.2 Glucose tolerance and insulin sensitivity

To further test our hypothesis, we performed IPGTT and IPIST post-treatment. CR monotherapy did not show an improvement in glucose tolerance and insulin sensitivity. GABA (*p*<0.05) and CR+GABA (*p*<0.01) treated groups showed a significant drop in the blood glucose levels by 120 mins as compared to HFD+STZ group (Figure 2A). The AUC plot showed a significant increase in glucose tolerance in GABA and CR+GABA treated groups (*p*<0.05) (Figure 2B). CR+GABA (*p*<0.01) treated group showed improved insulin sensitivity as the blood glucose levels were reduced to normoglycemia by 120 mins as compared to HFD+STZ group (Figure 2C & D).

**Figure 2.**
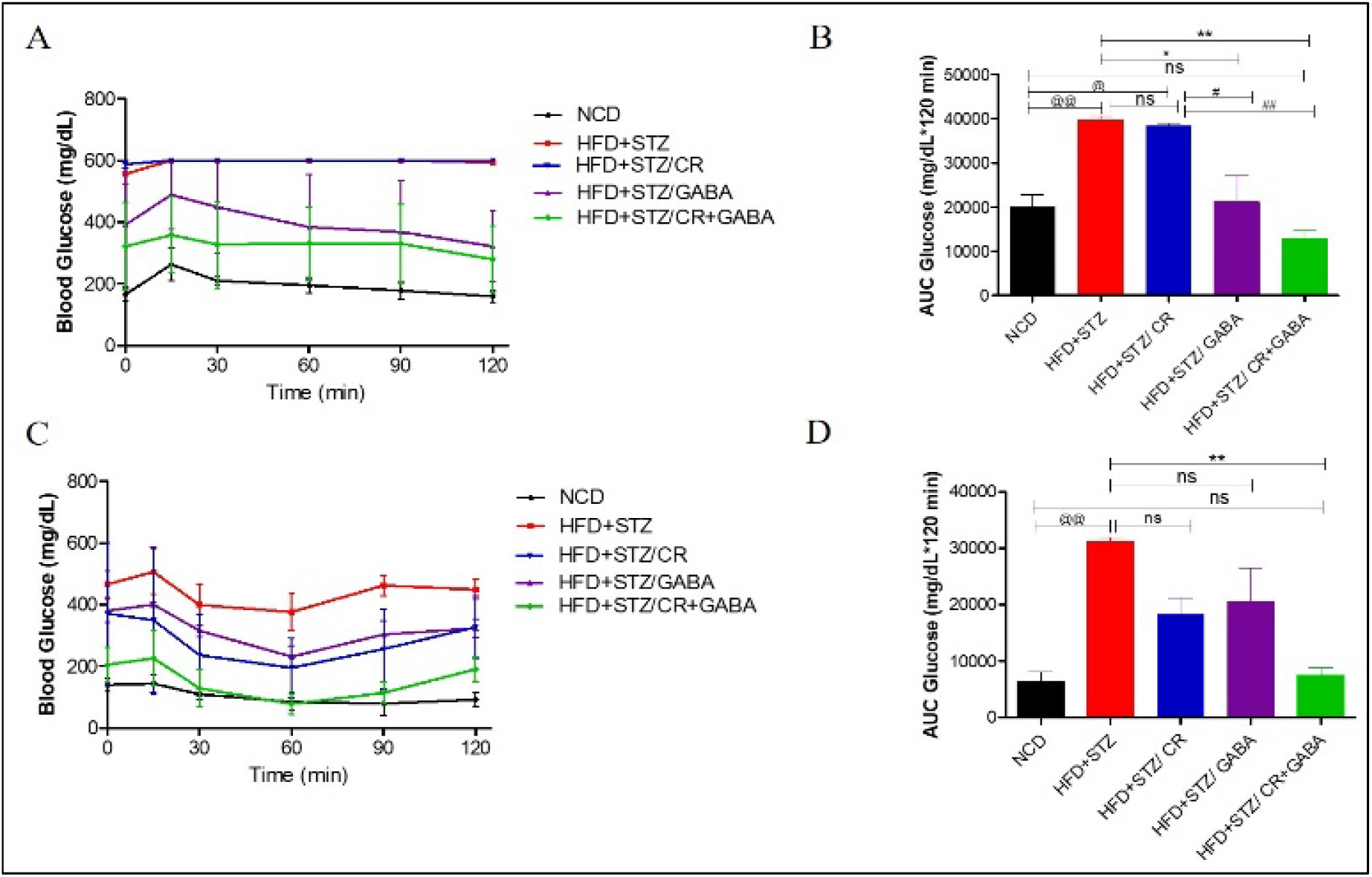
IGTT and IPIST. A) Blood glucose levels in the GABA and CR+GABA groups were significantly lower than CR and HFD+STZ mice at 60, 90 and 120 mins of glucose administration. B) AUC 0–120 curve indicated improved glucose tolerance in GABA and CR+GABA treated groups compared to HFD+STZ group. C) Blood glucose levels in CR+GABA was significantly lower than in CR, GABA and HFD+STZ groups at 60, 90 and 120 mins of insulin administration. D) AUC 0–120 curve in mice treated with CR+GABA combination was significantly lower than in HFD+STZ group. (ns=non-significant, \**p*<0.05, \*\**p*<0.01, ***p<0.001 vs HFD+STZ; ^@^*p*<0.05, ^@@^*p* <0.01 vs NCD; ^#^*p*<0.05, ^##^*p* <0.01 vs CR n=4-5/ group).

#### 3.2.4 Insulin and C-peptide levels

A significant increase in insulin and c-peptide levels along with improved insulin/c-peptide ratio was observed in GABA & CR+GABA (*p*<0.05) treated groups as compared to HFD+STZ group (Figure 4 A-C). No significant difference was observed in CR group.

**Figure 3.**
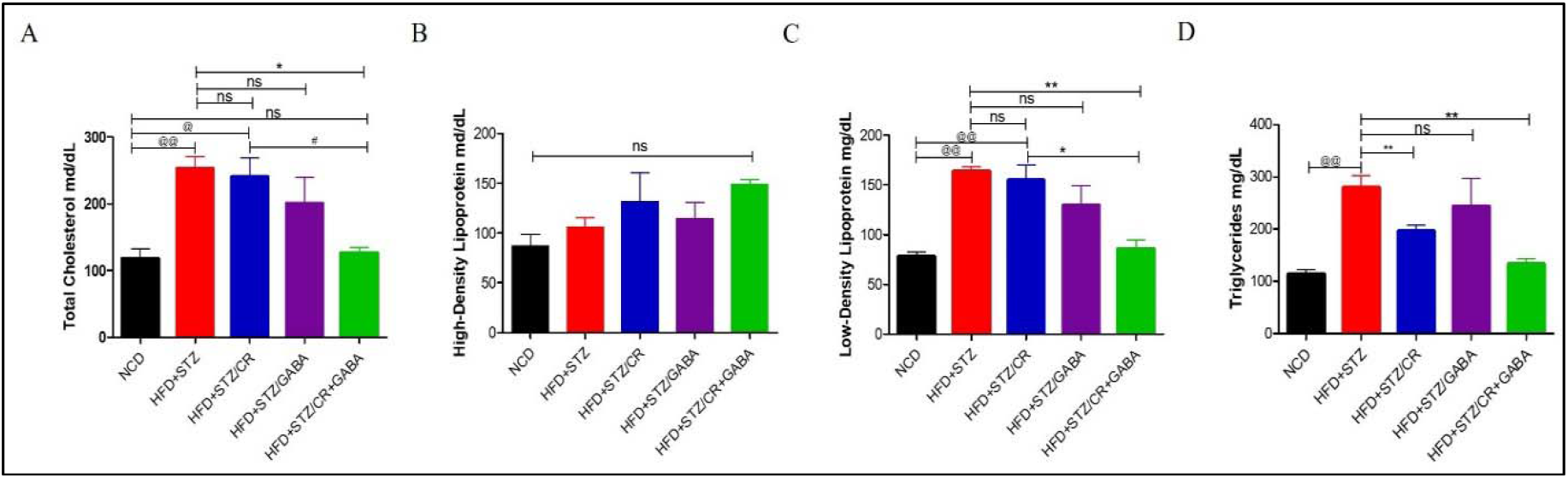
Plasma lipid levels. A-D) TC, TG, and LDL levels were reduced in CR+GABA group as compared to HFD+STZ group while no difference was observed in the rest of the treatment groups (ns=non-significant, \**p*<0.05, \*\**p*<0.01, vs HFD+STZ; ^@^*p*<0.05, ^@@^*p*<0.01 vs NCD; ^#^*p*<0.05, vs CR n=4-5/ group).

**Figure 4.**
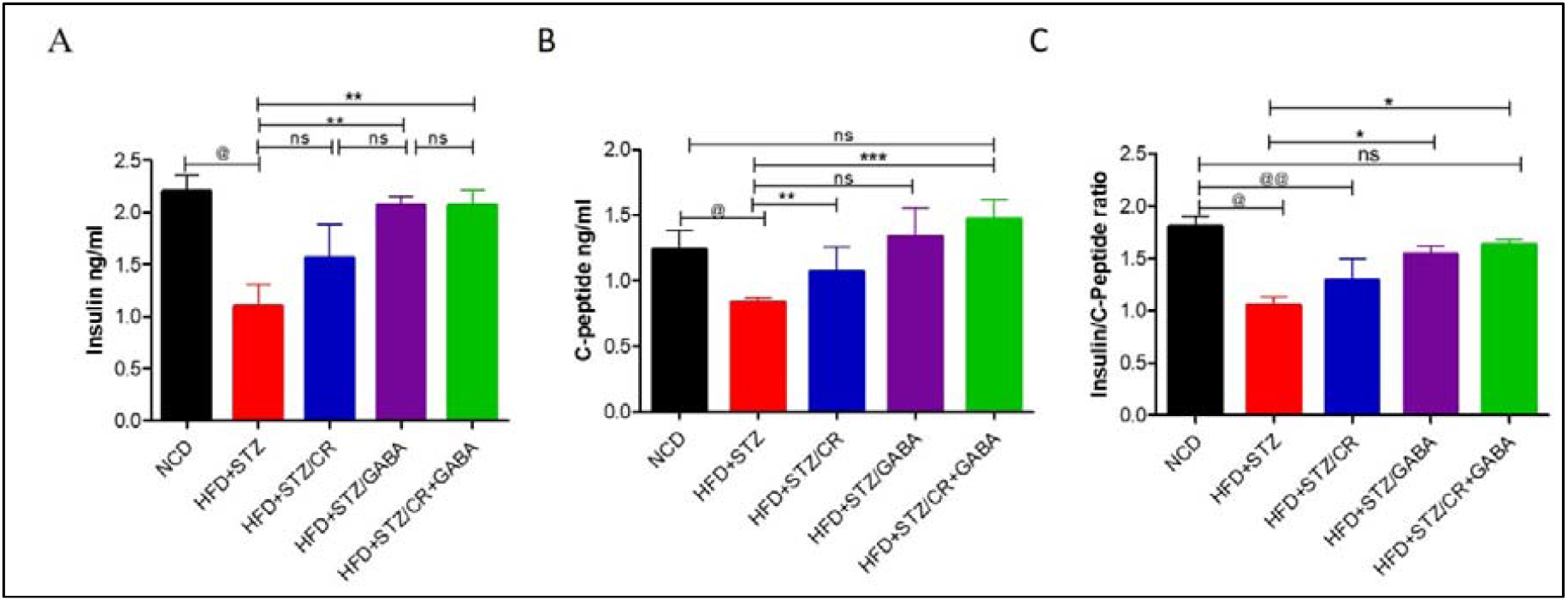
Plasma insulin, c-peptide levels and insulin/c-peptide ratio. (A & B) Increased insulin and c-peptide levels were observed in GABA & CR+GABA treated groups as compared to HFD+STZ. C) A significant increase in the insulin/c-peptide ratio was observed in GABA & CR+GABA treated groups as compared to HFD+STZ group (ns=non-significant, \**p*<0.05, \*\**p*<0.01, \*\*\**p*<0.001 vs HFD+STZ; ^@^*p*<0.05, ^@@^*p* <0.01 vs NCD; n=4-5/ group).

### 3.2 Gene expression profiling

GABA treated group showed a significant decrease (*p*<0.05) in the expression of *phosphoenolpyruvate carboxykinase* (*PEPCK-*gluconeogenesis), glucose-6 phosphatase (*G6Pase*), and *glycogen phosphorylase* (*GP*-glycogenolysis) in the liver (Figure 5). However, there was no significant difference in the expression of *adipose triglyceride lipase* (*ATGL*-lipolysis) and *acetyl-CoA carboxylase-1* (*ACC-1*-lipogenesis) in the adipose tissue (*p*>0.05) (Figure 6). Also, there was no significant difference observed in *sirtuin-1* (*SIRT-1*), *PPARG coactivator 1-*α (*PGC-1*α), and *transcription factor A mitochondrial* (*TFAM*-mitochondrial biogenesis) transcript levels in the skeletal muscle (*p*>0.05) (Figure 7). CR fed mice showed a significant decrease (*p*<0.05) in the expression of *G6Pase, PEPCK* (gluconeogenesis), *GLUT2* and *GP* (glycogenolysis) in the liver (Figure 5). However, there was no significant difference observed in the expression of *ATGL* and *ACC-1* in adipose tissue of CR fed mice (*p*>0.05) (Figure 6). Further, CR fed mice showed a significant increase (*p*<0.05) in *PGC-1*α transcript levels in skeletal muscle (Figure 7). CR+GABA treated group showed a significant decrease (*p*<0.05) in the expression of *G6Pase, PEPCK* (gluconeogenesis), *GLUT2* and *GP* (glycogenolysis) genes, and a significant increase (*p*<0.05) in *glucokinase* (glycolysis) expression (Figure 5). Interestingly, lipogenesis got up-regulated as marked by the increased *ACC-1* expression. Consequently, as a compensatory mechanism, lipolysis was also up-regulated, as observed by the increased *ATGL* expression (*p*<0.05) (Figure 6). CR+GABA treated group also showed a significant increase (*p*<0.05) in *SIRT-1, PGC-1*α and *TFAM* transcript levels in skeletal muscle (Figure 7).

**Figure 5.**
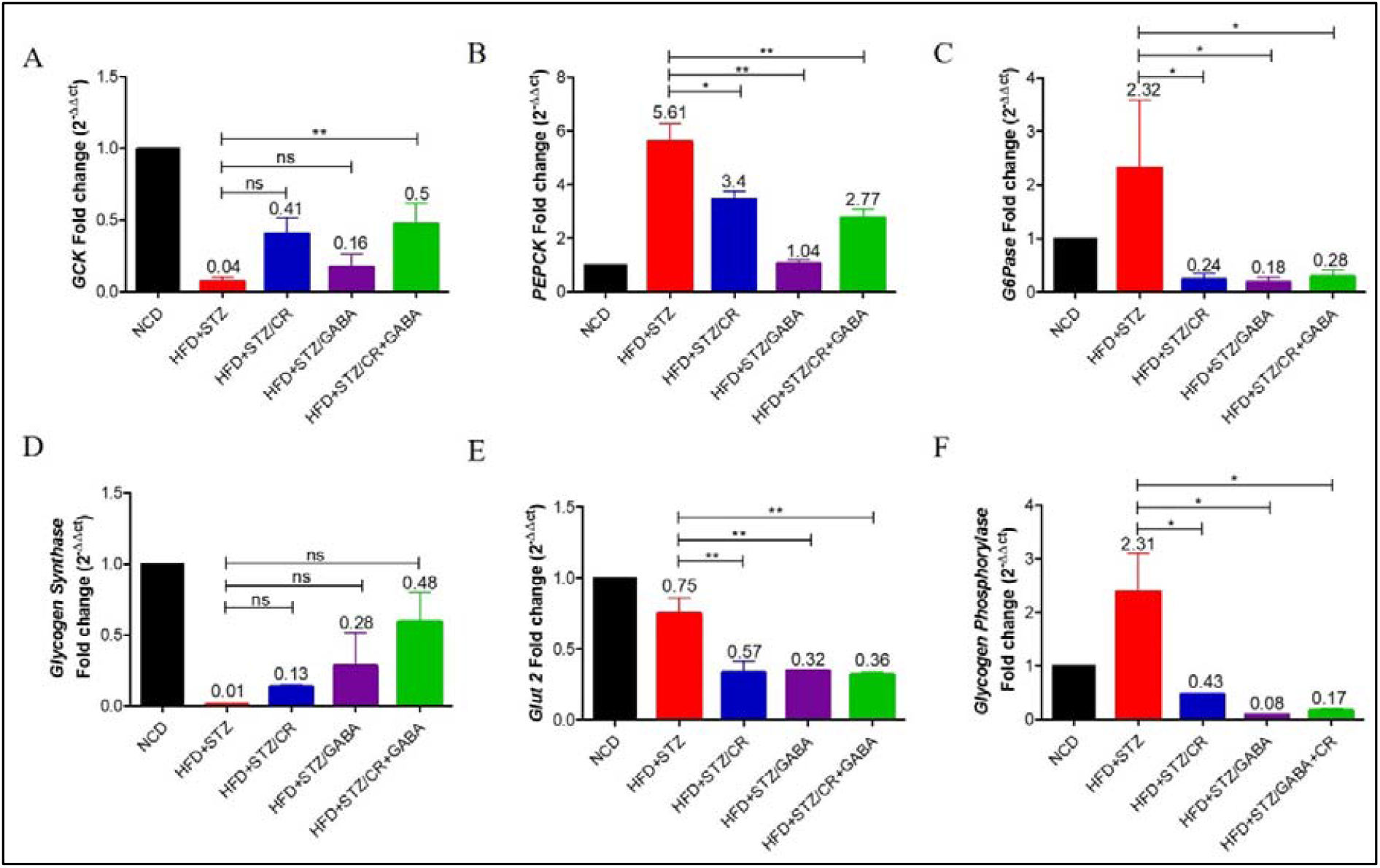
Transcript levels of glucoregulatory enzymes in the liver: **A. *GCK* fold change.** After normalisation with GAPDH expression, a 0.5-fold increase in *GCK* transcript levels was observed in the CR+GABA group as compared to the HFD+STZ group. **B & C. Fold change in *PEPCK* & *G6Pase***. GABA, CR and CR+GABA showed a significant 1-fold, 3-fold and 2.7 fold decrease in *PEPCK*. GABA, CR, and CR+GABA showed a significant 0.18-fold, 0.24-fold and 0.28-fold decrease in G6Pase expression compared to the HFD+STZ group. **D-E**.***Glycogen synthase, GLUT2* fold change**. No significant change was observed in *glycogen synthase*, whereas *GLUT2* expression decreased in all the treated groups compared to the HFD+STZ group. **F. Fold change in *Glycogen phosphorylase***. A significant 0.08-fold, 0.43-fold and 0.17-fold reduction in *glycogen phosphorylase* expression was observed in the treatment groups compared to the HFD+STZ group. (ns=non-significant, \**p*<0.05, \*\**p*<0.01; \*\*\**p*<0.001, n=4-5/group).

**Figure 6.**
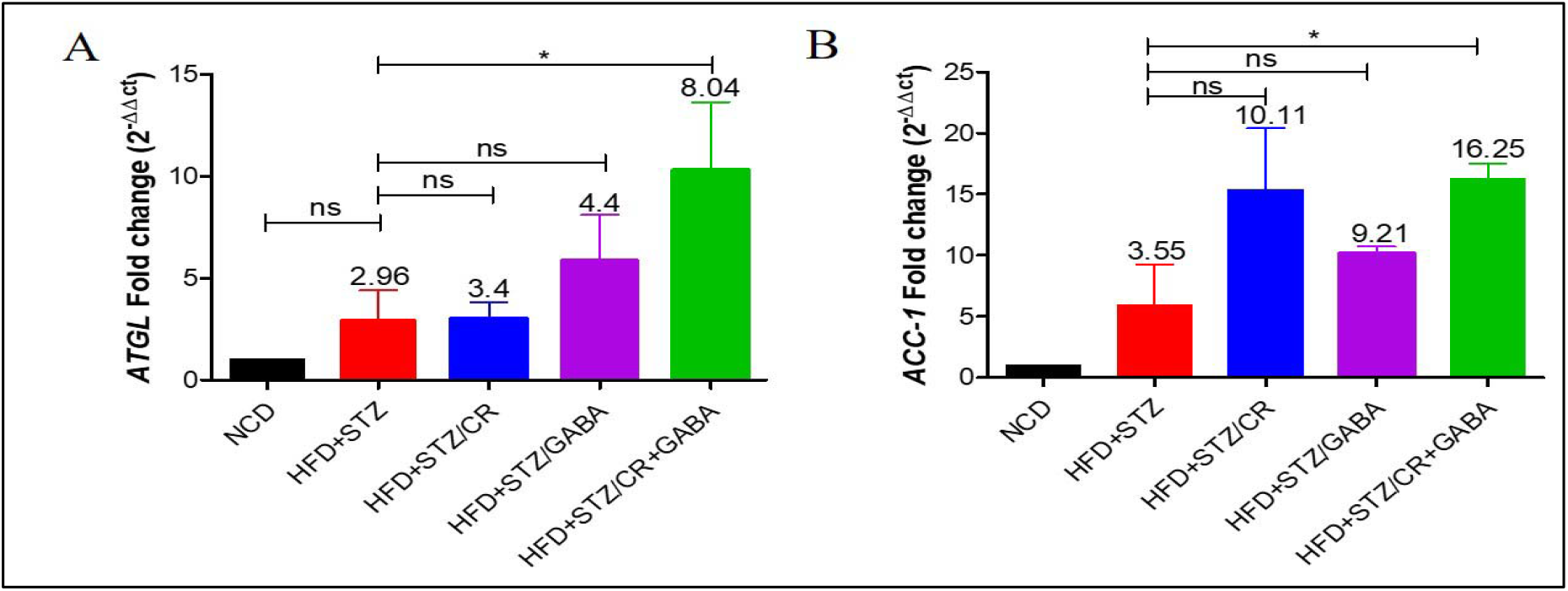
Transcript levels of lipid metabolism markers in the adipose tissue. A) Fold change in ***ATGL* mRNA**. A significant 8-fold increase in *ATGL* expression could be seen in the CR+GABA combination group as compared to the HFD+STZ group. B) Fold change in ***ACC-1* mRNA**. A significant 16-fold increase in *ACC-1* expression was seen in the CR+GABA group compared to the HFD+STZ group. (\**p*<0.05, ns=non-significant; *p*>0.05) (n=4-5/group).

**Figure 7.**
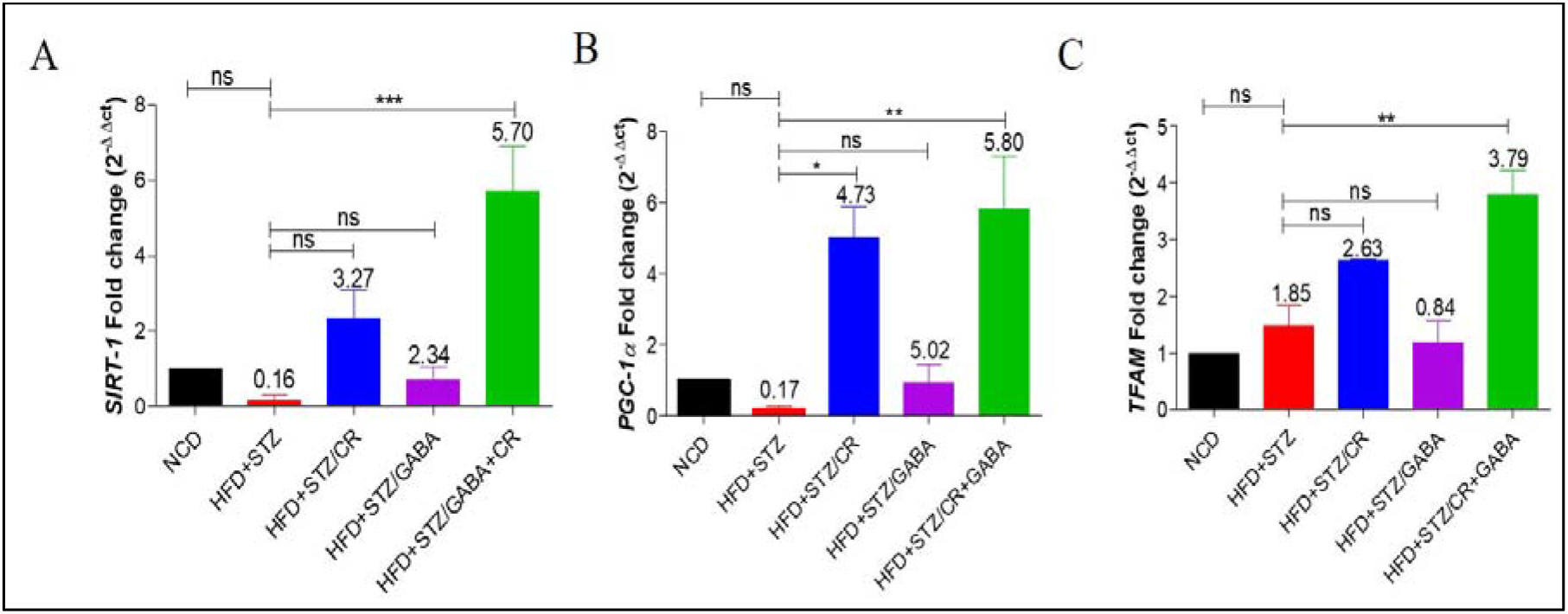
Transcript levels of mitochondrial biogenesis markers in the skeletal muscle. A) Fold change in ***SIRT-1* mRNA**. After normalisation with GAPDH expression, 5.7 fold increase in *SIRT-1* expression was observed in the CR+GABA treated group as compared to HFD+STZ group. B) ***PGC-1***α **mRNA fold change**. A significant increase in *PGC-1* expression was observed in CR (4.7-fold) and CR+GABA (5.8-fold) treated groups as compared to HFD+STZ group. C) ***TFAM* mRNA fold change**. A significant increase in *TFAM* expression (3.7-fold) was observed in the CR+GABA treated group as compared to HFD+STZ group. (*p<0.05, **p<0.01, ns=non-significant; p>0.05) (n=4-5/group).

### 3.4 Oxygen consumption rate

The rate of oxygen consumption is a significant indicator of mitochondrial activity. GABA group showed no significant difference in the ETC complex activities in skeletal muscle (*p*>0.05). CR fed mice showed a significant increase (*p*<0.05) in OCR by ETC complex I compared to the HFD+STZ group, indicating improved complex I activity in skeletal muscle. The combination therapy group showed significant OCR by ETC complexes I-III as compared to the HFD+STZ group indicating improved complex activities in skeletal muscle (*p*<0.05) (Figure 8).

**Figure 8.**
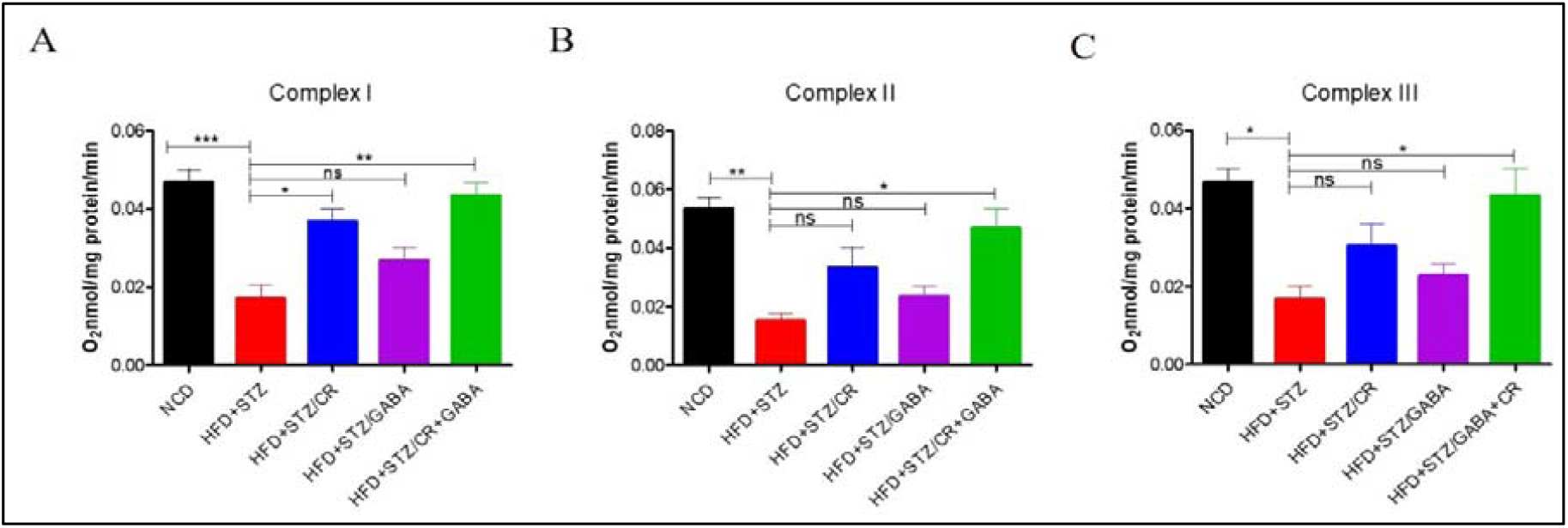
Estimation of OCR in the skeletal muscle. The activities of complexes I-III were significantly reduced in the HFD+STZ group. A) CR and CR+GABA treated groups showed a significant increase in complex I as compared to HFD+STZ. B-C) CR+GABA treated group showed a significant increase in complexes II and III activities compared to the HFD+STZ group. (ns=non-significant, \**p*<0.05, \*\**p*<0.01, \*\*\**p*<0.001, n=4-5 group).

### 3.5 Pancreatic β-cell regeneration and apoptosis

β-cell proliferation and apoptosis were assessed by IHC microscopy. The GABA treated group showed a significant increase in β-cell proliferation [BrdU/Insulin co-positive cells] (Figure 9) as compared to NCD (*p*<0.01) and HFD+STZ treated groups (*p*<0.001). A similar trend was observed for neogenesis [PDX-1/NGN-3/Insulin co-positive cells] (Figure 10) as compared to NCD (*p*<0.001) and HFD+STZ treated groups (*p*<0.001). Also, GABA treated group showed a significant reduction (*p*<0.01) in β-cell apoptosis [TUNEL positive cells] (Figure 12). However, the CR monotherapy group showed no β-cell regeneration, and there was no improvement in β-cell apoptosis (*p*>0.05) (Figure 9-12). The CR+GABA treated group showed a significant increase as compared to NCD and HFD+STZ in β-cell proliferation (*p*<0.001; *p*<0.001) (Figure 9) and neogenesis (*p*<0.001; *p*<0.001) (Figure 10). The CR+GABA treated group showed significantly reduced β-cell apoptosis as compared to HFD+STZ (*p*<0.001) (Figure 12). The GABA and CR monotherapies, along with the GABA+CR group, did not show transdifferentiation (ARX/PAX4/Insulin co-positive cells) (*p*>0.05) (Figure 11). Further, to validate β-cell apoptosis we used AIF as a marker. However, no AIF translocation was observed in any of the groups (Figure 13).

**Figure 9.**
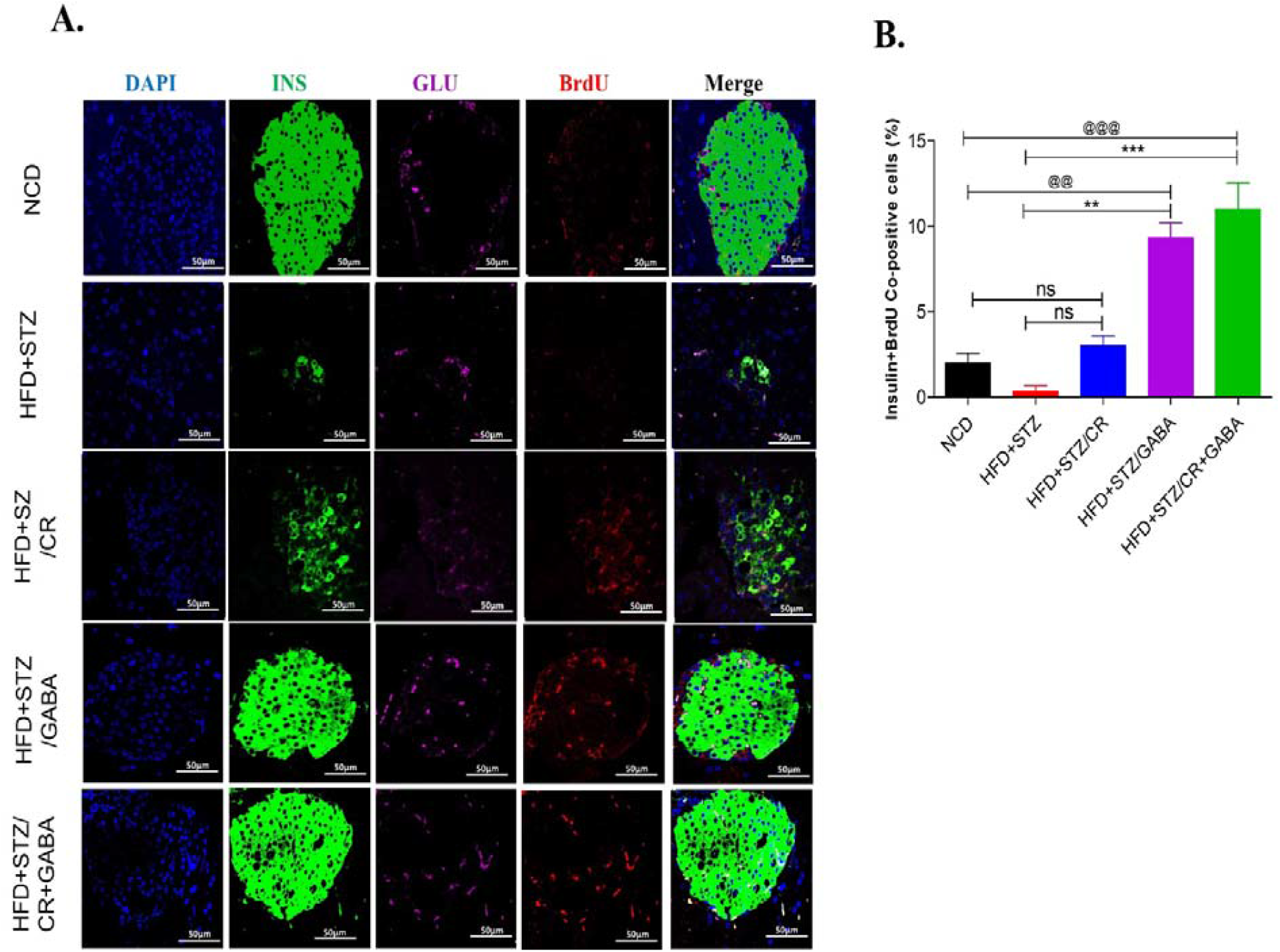
Assessment of pancreatic β-cell proliferation. (A) Representative immunofluorescence images of pancreatic islets showing insulin (green), glucagon (magenta) and BrdU (red) cells in all the groups. (B) Percentage of Insulin and BrdU co-positive cells. GABA and CR+GABA treated groups showed a significant increase in β-cell proliferation as compared to the HFD+STZ group. However, no significant difference was observed in the CR group. (ns=non-significant, \*\**p*<0.01, \*\*\**p*<0.001 vs HFD+STZ; ^@@^*p*<0.01, ^@@@^*p*<0.001 vs NCD; n=3/ group).

**Figure 10.**
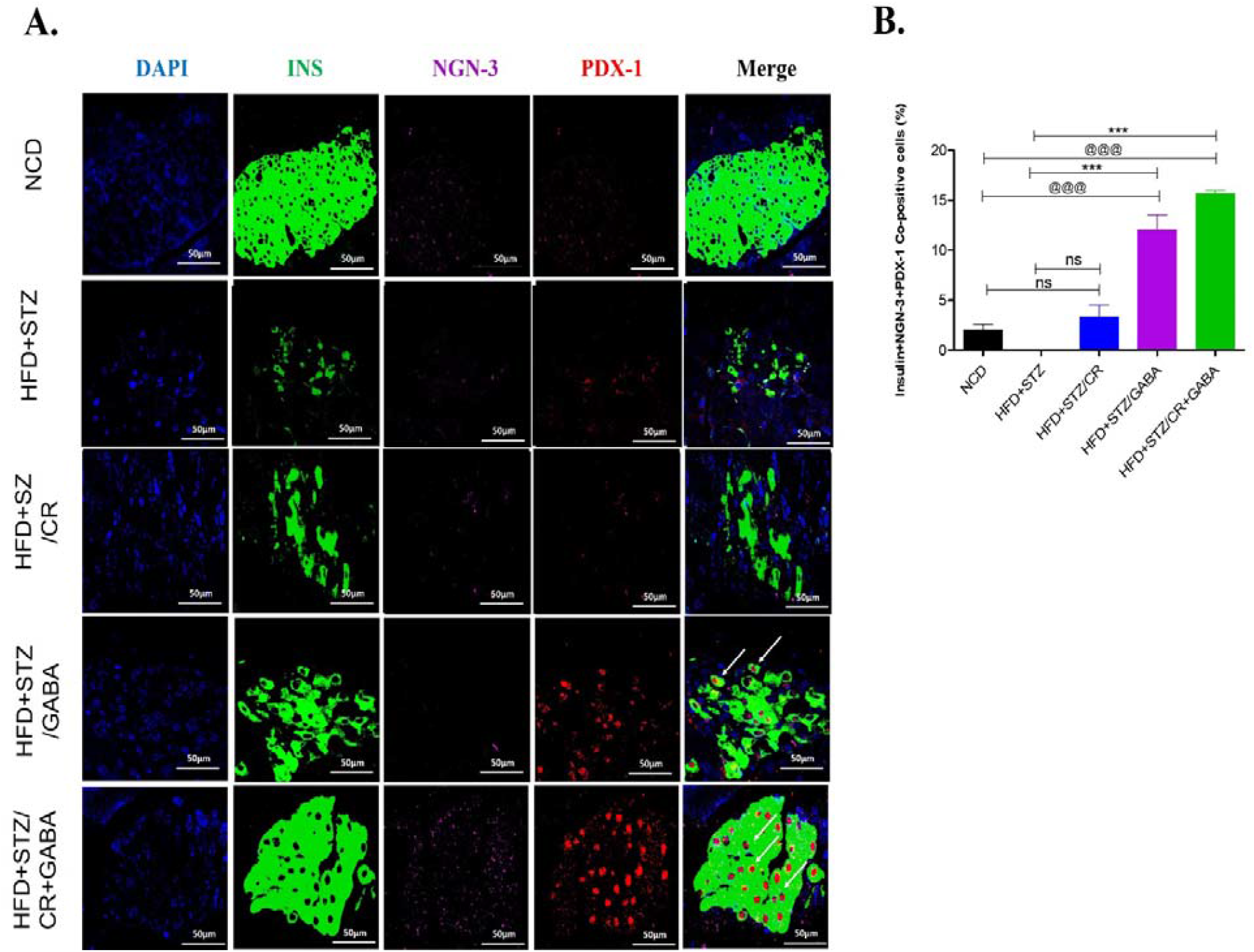
Assessment of pancreatic β-cell neogenesis. **(A)** Representative immunofluorescence images of pancreatic islets showing insulin (green), NGN-3 (magenta) and PDX-1 (red) cells in all the groups. (B) Percent of Insulin, NGN-3 and PDX-1 co-positive cells. GABA and CR+GABA treated groups showed a significant increase in β-cell neogenesis as compared to the HFD+STZ group. However, no significant difference was observed in the CR group. (ns=non-significant, \*\*\**p*<0.001 vs HFD+STZ; ^@@@^*p*<0.001 vs NCD; n=3/ group).

**Figure 11.**
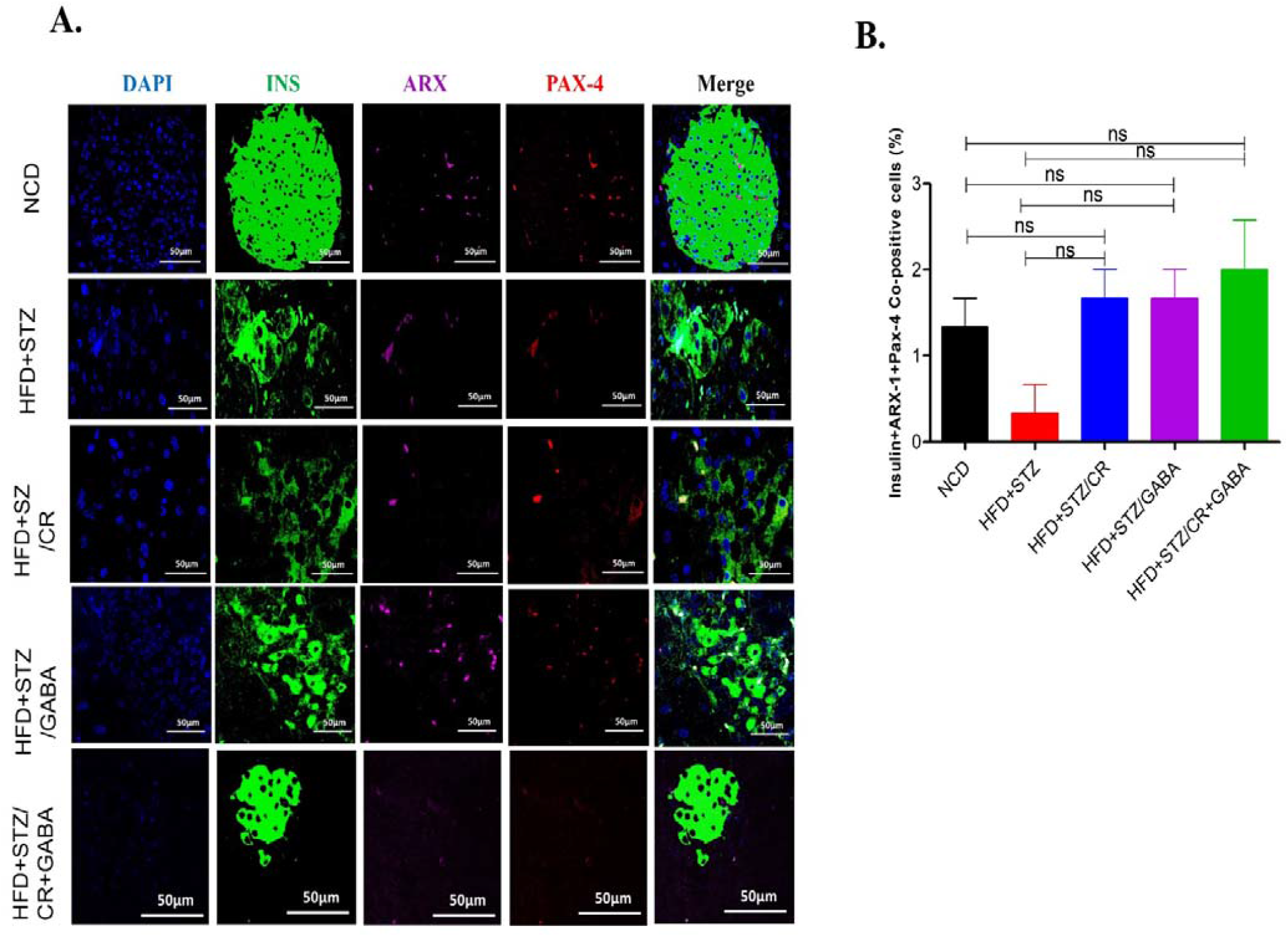
Assessment of pancreatic β-cell transdifferentiation. **(A)** Representative immunofluorescence images of pancreatic islets showing insulin (green), ARX-1 (magenta) and PAX-4 (red) cells in all the groups. (B) Percent of Insulin, ARX, and PAX-4 co-positive cells. β-cell transdifferentiation was not seen in any of the treated groups as compared to the HFD+STZ group. (ns=non-significant; n=3/ group).

**Figure 12.**
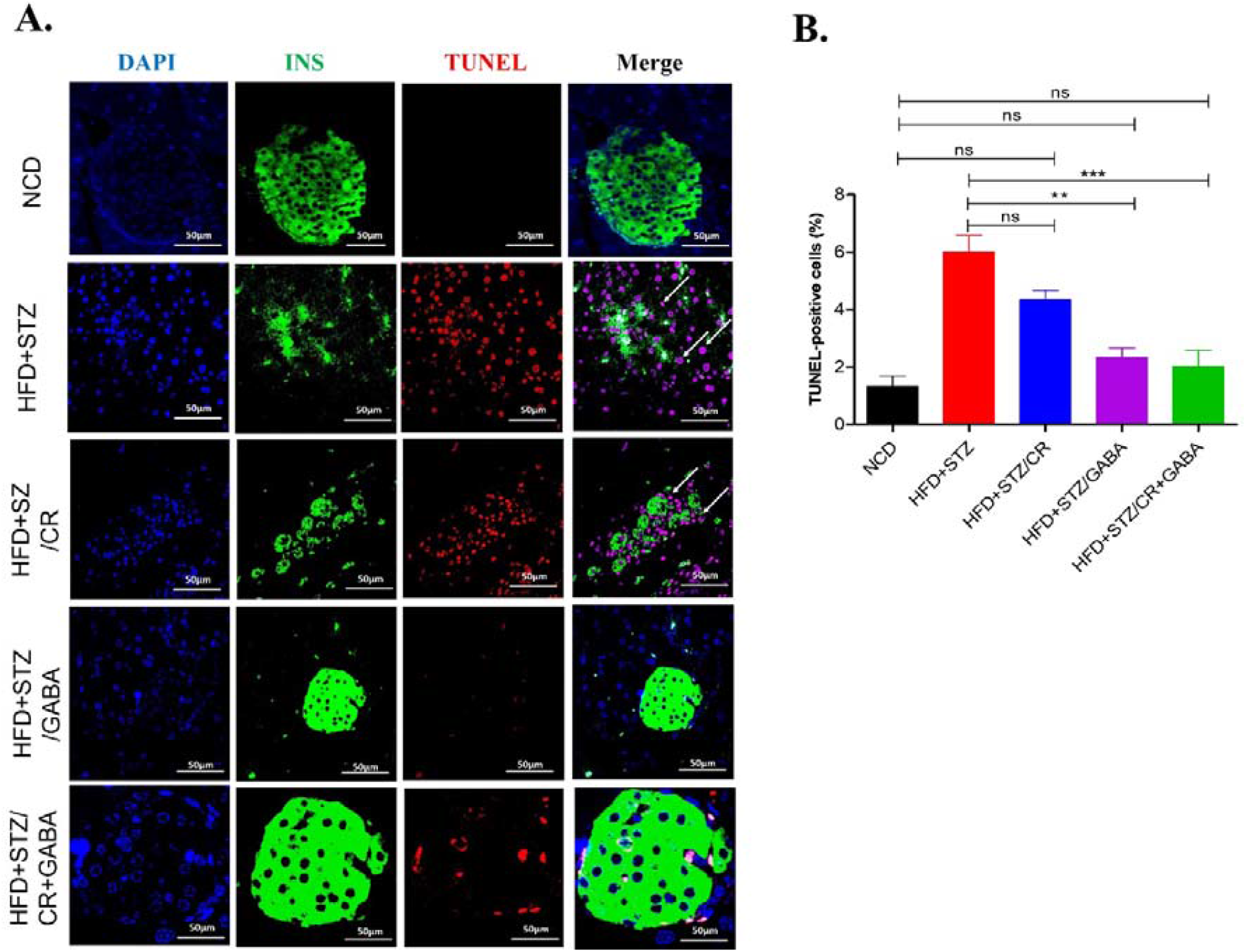
Assessment of pancreatic β-cell apoptosis: (A) Representative immunofluorescence images of pancreatic islets, showing insulin (green) and TUNEL (red) cells in all the groups. (B) Percent of Insulin and TUNEL co-positive cells. A significant reduction in β-cell apoptosis was observed in GABA and CR+GABA groups compared to the HFD+STZ group. (ns=non-significant, \*\**p*<0.01, \*\*\**p*<0.001 vs HFD+STZ; n=3/group).

**Figure 13.**
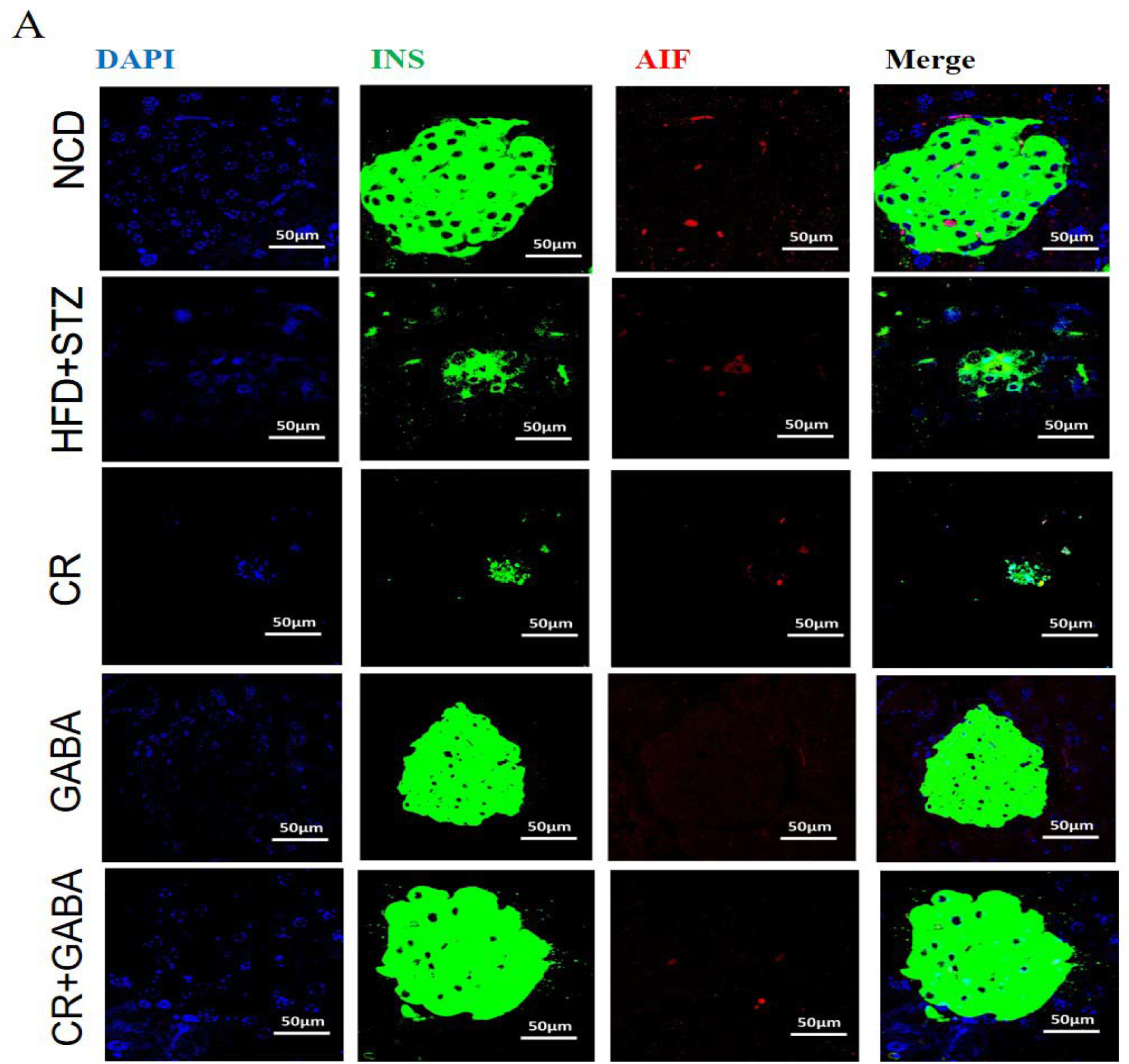
Assessment of AIF translocation: (A) Representative immunofluorescence images of pancreatic islets, showing insulin (green) and AIF (red) cells in all the groups. No significant translocation of AIF was observed in any of the groups. (n=3/group).

## 4. Discussion

The management of T2D involves two main approaches: i) increasing insulin secretion from the β-cells and ii) increasing insulin-mediated glucose uptake by the peripheral tissues. Patients develop tolerance against the existing line of treatment within a few years, which poses a challenge for developing new medicines. These drugs also lead to many side-effects [1]. Thus, combination therapies involving naturally existing biomolecules are the way forward to overcome the complications of T2D. Therefore, the present study was designed to determine the therapeutic potential of a combination of CR and GABA in the T2D mouse model.

CR is a potential dietary intervention emphasizing caloric management in terms of fats, carbohydrates and proteins with sufficient vitamins and minerals. CR promotes insulin sensitivity, mitochondrial biogenesis and functionality [13, 15]. Mice fed with 30-40% of the ad libitum calorie deficit diet showed increased life expectancy, robustness against toxicity and stress [27]. Further data on similar lines showed CR’s role in reducing body weight and lipid profile markers when the therapy was prolonged [28]. Several reports have explained CR’s role as an activator of the SIRT1-PGC-1α signalling pathway at the transcriptional level [30]. CR modulates lipid metabolism via the SIRT-1 signalling pathway and reduces intracellular diacylglycerol species [29]. In the liver, *PGC-1*α may regulate the gluconeogenic pathway; however, the molecular mechanism remains unknown [31].

Interestingly, we have also seen similar observations of action at multiple sites with CR monotherapy reducing body weight and triglyceride levels and improving insulin sensitivity besides reducing gluconeogenesis and glycogenolysis. Civitarese *et al*. studied the molecular mechanism of CR in mitochondrial functionality. They explained an increased mRNA expression of the genes involved in mitochondrial biogenesis (viz. *SIRT1, PGC-1*α and *TFAM*), with *SIRT1* being the direct regulator of the pathway [32]. Increased mitochondrial biogenesis and oxygen consumption rate by ETC complex I in our CR monotherapy group find agreement with their findings.

GABA has a proven role in islet□cell hormone homeostasis, preserving the β□cell mass, suppressing immune reactions and consequent apoptosis [19, 33]. The food sources of GABA are potatoes, tomatoes and brown rice, while microorganisms like *E. coli, Lactococcus lactis* produce it [35]. In the present study, GABA monotherapy effectively reduces FBG levels, improves glucose tolerance, increases insulin and c-peptide levels and reduces gluconeogenesis and glycogenolysis. GABAnergic system functions in many tissues, including the peripheral tissues. In the liver, GABAAR regulates PI3/Akt activities that maintain hepatocyte survival and *PGC-1*α expression. Thus, GABA is one of the multiple factors responsible for regulating hepatic glucose metabolism, i.e. gluconeogenic and glycogenolytic pathways [36]. Fibroblast growth factor 21 (FGF-21) predominantly expressed in the liver under normal condition also induces *PGC-1*α expression. It leads to the regulation of lipolysis in the adipose tissue forming liver-adipose tissue crosstalk [37]. The positive effect of GABA treatment stands documented in T1D and T2D murine models. GABA therapy protects NOD animals from diabetes, and a similar effect is reported in various *in-vivo* models [20, 17]. Also, GABA regulates cytokine secretion from human PBMCs and suppresses β-cell□reactive CD8+ CTLs in T1D models [19, 17], proposing the role of GABA as an immunosuppressant. GABA reportedly acts as an inducer of α-to-β-like cell conversion *in-vivo* upon prolonged exposure in the STZ-induced mouse model [19]. Although no β-cell transdifferentiation is seen in our GABA-treated group, a significant increase in β-cell proliferation and neogenesis and reduced β-cell apoptosis were observed. The individual effects of CR and GABA on the multiple pathways and tissues diminishe the adverse actions of T2D pathophysiology when given in combination. Our results suggest that the combination treatment improves glucose homeostasis by increasing insulin sensitivity and glucose tolerance. CR enhances insulin sensitivity by improving the mitochondrial function at two levels, elevating the expression of *SIRT1, PGC-1*α and *TFAM* and increasing mitochondrial complex I-III activities. GABA therapy boosts insulin levels by inducing β-cell proliferation and neogenesis and contributes to the existing depositary. GABA also enhances the metabolic profile by reducing BW, FBG levels, triglycerides, total cholesterol, and LDL levels. As mentioned earlier, the liver-adipose tissue crosstalk regulates these pathways at the transcriptional level. CR+GABA treated group shows a significant decrease in the expression of *G6Pase* and *PEPCK* (gluconeogenesis), *GLUT2* and *G6Pase* (glycogenolysis), and an increase in *glucokinase* (glycolysis). Interestingly, lipogenesis seems up-regulated as marked by the increased *ACC* expression, and consequently, lipolysis also gets up-regulated as a compensatory mechanism as observed by the increased *ATGL* expression. CR+GABA treated group showed a significant increase in β-cell proliferation, neogenesis and reduced β-cell apoptosis. We did not observe AIF translocation for β-cell apoptosis, a marker for caspase-independent cell death. Increased oxidative stress induced by STZ and low levels of antioxidant enzymes in the pancreatic cells activate caspase-3 and caspase-9 pathway to mediate β-cell apoptosis [38-40]. Hence, the HFD+STZ model mimics caspase-dependent β-cell apoptosis. However, further validation is required for caspase mediated cell death pathways.

However, a few limitations of this study are as follows. First, we used CR as one of our therapies and monitored parameters at the transcriptional level in different peripheral tissues. As CR is first-line therapy, it is ideal for assessing energy expenditure parameters. Second, our group has not performed cell lineage tracing studies to establish GABA’s role in β-cell regeneration. Thus, extensive *in vitro* and *in vivo* investigations are needed to address the above burning question and whether the effect was due to the two different signalling cascades additive stimulus or an undefined underlying mechanism. Thus, the current study suggests that combination therapy with CR and GABA leads to amelioration of T2D pathophysiology in the T2D mouse model (Figure 14).

**Figure 14.**
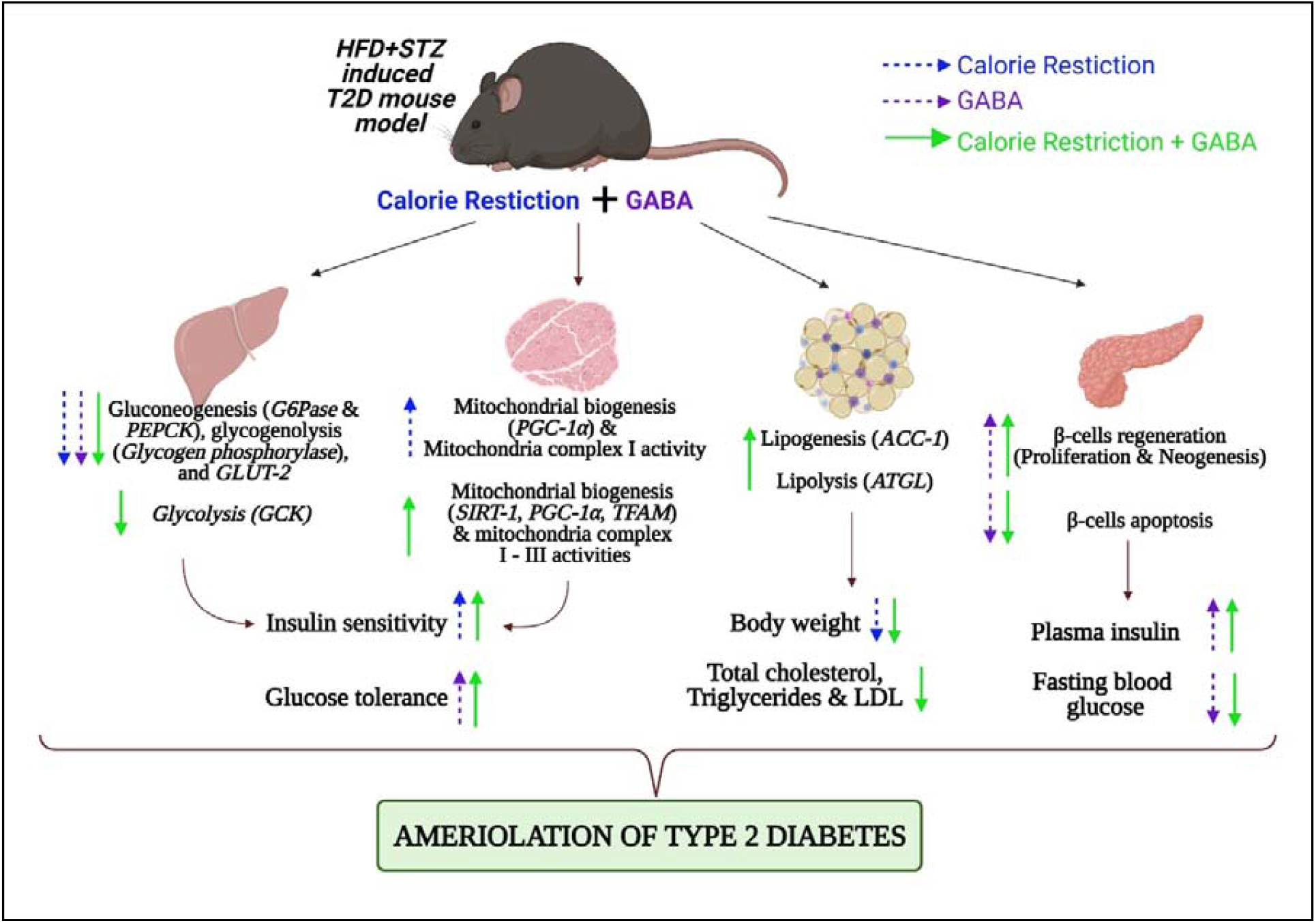
The effect of GABA, CR and CR+GABA (combination therapy) on amelioration of T2D pathophysiology in HFD+STZ induced T2D mouse model. T2D pathophysiology is characterized by insulin resistance and β-cell loss. GABA monotherapy shows reduced FBG levels, improved insulin sensitivity and glucose tolerance, increased insulin and c-peptide levels and decreased gluconeogenesis and glycogenolysis. The GABA treated group also shows a significant increase in β-cell proliferation and neogenesis with significantly reduced β-cell apoptosis. CR diet-fed mice show reduced body weight and triglycerides levels, along with improved insulin sensitivity, reduced gluconeogenesis and glycogenolysis. These mice also show elevated expression of mitochondrial biogenesis markers and oxygen consumption rate by ETC complex I-III compared to the HFD+STZ group. CR diet-fed mice show no β-cell regeneration and no improvement in β-cell apoptosis compared to the HFD+STZ group. GABA+CR treated group shows improved glucose homeostasis by increasing insulin sensitivity and glucose tolerance, enhancing the transcript levels of key markers of glucoregulatory enzymes and lipid metabolism; increasing mitochondrial biogenesis and ETC complex activities. Further, the combination treatment promotes β-cell regeneration and reduces β-cell apoptosis compared to the HFD+STZ group.

## 5. Conclusion

Our study suggests that CR in combination with GABA therapy could help overcome T2D pathophysiology by their action at multiple sites. CR regulates glucose and lipid metabolism, mitochondrial biogenesis, ETC complex activities, promotes β-cell regeneration and reduces β-cell apoptosis. CR treatment seems part of a first-line therapy that can be used with other drugs like DPP-IV inhibitors or metformin. Moreover, a CR-mimetic, for instance, resveratrol, is available commercially to easily overcome the challenge of strict dietary regimen and protocols. GABA monotherapy works on β-cell regeneration, readily available from natural sources like spinach and tomatoes. An edible vegetable extract can be formulated for diabetic patients without the manifestation of significant side effects. Thus, combination therapy can be useful for effective action at multiple sites and mechanisms to combat T2D pathophysiology. This study needs further investigation at pre-clinical and clinical trial levels to evaluate the commercial viability.

## Abbreviations

T2D: Type 2 Diabetes
FBG: Fasting Blood Glucose
TC: Total Cholesterol
HDL: High Density Lipoprotein
TG: Triglycerides
LDL: Low Density Lipoprotein
BMI: Body Mass Index
HFD: High Fat Diet
CR: Calorie Restriction
STZ: Streptozocin
GABA: γ-Aminobutyric Acid

## Acknowledgments

We thank Dr. Ansarullah, Institute of Diabetes and Regeneration Research, Helmholtz Center, Munich, Neuherberg, Germany for the project idea. We thank Dr. Deepak Sharma, Scientist F, BARC, Mumbai, for providing the confocal facilities. NR thanks University Grants Commission-National Fellowship for higher education for ST students, New Delhi, India, for awarding SRF. RP thanks Council for Scientific and Industrial Research, New Delhi, India, for awarding SRF. Images are created with BioRender.com

## Author Contributions

RB, NR, AVR and RBS conceived the idea. NR designed and performed the experiments; did data acquisition and data analysis; and wrote the original manuscript draft. NP performed a few experiments and data acquisition. SP and RP reviewed and edited the manuscript. RB and AVR contributed to the critical revision and approval of the manuscript.

## References

1. Rathwa N, Patel R, Palit SP, Parmar N, Rana S, Ansari MI, Ramachandran AV, Begum β-Cell Replenishment: Possible Curative Approaches for Diabetes Mellitus. Nutrition, Metabolism and Cardiovascular Diseases. 2020 Aug 11.

2. Dwivedi M, Laddha NC, Imran M, Ansarullah Ramachandran AV, Misra A, Yadav M, Begum R. ACE gene I/D polymorphism in type 2 diabetes: the Gujarat population. The British Journal of Diabetes & Vascular Disease. 2011 May;11(3):153–4.

3. Patel R, Dwivedi M, Mansuri MS, Laddha NC, Thakker A, Ramachandran AV, Begum Association of neuropeptide-Y (NPY) and interleukin-1beta (IL1B), genotype-phenotype correlation and plasma lipids with Type-II diabetes. PloS one. 2016 Oct 17;11(10):e0164437.

4. Patel R, Rathwa N, Palit SP, Ramachandran AV, Begum R. Association of melatonin & MTNR1B variants with type 2 diabetes in Gujarat population. Biomedicine & Pharmacotherapy. 2018 Jul 1;103:429–34.

5. Patel R, Palit SP, Rathwa N, Ramachandran AV, Begum R. Genetic variants of tumor necrosis factor-α and its levels: A correlation with dyslipidemia and type 2 diabetes susceptibility. Clinical nutrition. 2019 Jun 1;38(3):1414–22.

6. Palit SP, Patel R, Jadeja SD, Rathwa N, Mahajan A, Ramachandran AV, Dhar MK, Sharma S, Begum R. A genetic analysis identifies a haplotype at adiponectin locus: Association with obesity and type 2 diabetes. Scientific reports. 2020 Feb 19;10(1):1.

7. Rathwa N, Patel R, Palit SP, Ramachandran AV, Begum R. Genetic variants of resistin and its plasma levels: Association with obesity and dyslipidemia related to type 2 diabetes susceptibility. Genomics. 2019 Jul 1;111(4):980–5.

8. Rathwa N, Parmar N, Palit SP, Patel R, Ramachandran AV, Begum R. Intron specific polymorphic site of vaspin gene along with vaspin circulatory levels can influence pathophysiology of type 2 diabetes. Life Sciences. 2020 Feb 15;243:117285.

9. Rathwa N, Patel R, Palit SP, Jadeja SD, Narwaria M, Ramachandran AV, Begum R. Circulatory Omentin-1 levels but not genetic variants influence the pathophysiology of Type 2 diabetes. Cytokine. 2019 Jul 1;119:144–51.

10. Pramanik, S., Rathwa, N., Patel, R., Ramachandran, A. V., & Begum, R. (2018). Treatment avenues for type 2 diabetes and current perspectives on adipokines. Current diabetes reviews, 14 (3), 201–221.

11. Rathwa, N., Patel, R., Palit, S. P., Ramachandran, A. V., & Begum, R. (2018). Genetic variants of resistin and its plasma levels: association with obesity and dyslipidemia related to type 2 diabetes susceptibility. Genomics.

12. Rathwa, N., Patel, R., Palit, S. P., Jadeja, S. D., Narwaria, M., Ramachandran, A. V., & Begum, R. (2019). Circulatory Omentin-1 levels but not genetic variants influence the pathophysiology of Type 2 diabetes. Cytokine, 119, 144–151.

13. López□Lluch G, Navas P. Calorie restriction as an intervention in ageing. The Journal of physiology. 2016 Apr 15;594(8):2043–60.

14. Golbidi S, Daiber A, Korac B, Li H, Essop MF, Laher I. Health benefits of fasting and caloric restriction. Current diabetes reports. 2017 Dec 1;17(12):123.

15. Picca A, Lezza AM. Regulation of mitochondrial biogenesis through TFAM– mitochondrial DNA interactions: useful insights from aging and calorie restriction studies. Mitochondrion. 2015 Nov 1;25:67–75.

16. Dogan S, Ray A, Cleary MP. The influence of different calorie restriction protocols on serum pro-inflammatory cytokines, adipokines and IGF-I levels in female C57BL6 mice: short term and long term diet effects. Meta gene. 2017 Jun 1;12:22–32.

17. Soltani N, Qiu H, Aleksic M, Glinka Y, Zhao F, Liu R, Li Y, Zhang N, Chakrabarti R, Ng T, Jin T. GABA exerts protective and regenerative effects on islet beta cells and reverses diabetes. Proceedings of the National Academy of Sciences. 2011 Jul 12;108(28):11692–7.

18. Purwana I, Zheng J, Li X, Deurloo M, Son DO, Zhang Z, Liang C, Shen E, Tadkase A, Feng ZP, Li Y. GABA promotes human β-cell proliferation and modulates glucose homeostasis. Diabetes. 2014 Dec 1;63(12):4197–205.

19. Wang Q, Ren L, Wan Y, Prud’homme GJ. GABAergic regulation of pancreatic islet cells: Physiology and antidiabetic effects. Journal of cellular physiology. 2019 Jan 28.

20. Tian J, Dang H, Kaufman DL. Combining antigen-based therapy with GABA treatment synergistically prolongs survival of transplanted ß-cells in diabetic NOD mice. PloS one. 2011 Sep 22;6(9):e25337.

21. Ben-Othman N, Vieira A, Courtney M, Record F, Gjernes E, Avolio F, Hadzic B, Druelle N, Napolitano T, Navarro-Sanz S, Silvano S. Long-term GABA administration induces alpha cell-mediated beta-like cell neogenesis. Cell. 2017 Jan 12;168(1-2):73–85.

22. Huang P, Li S, Shao M, Qi Q, Zhao F, You J, Mao T, Li W, Yan Z, Liu Y. Calorie restriction and endurance exercise share potent anti-inflammatory function in adipose tissues in ameliorating diet-induced obesity and insulin resistance in mice. Nutrition & metabolism. 2010 Dec;7(1):1–9.

23. Fontana L, Partridge L. Promoting health and longevity through diet: from model organisms to humans. Cell. 2015 Mar 26;161(1):106–18.

24. Bansal P, Paul P, Mudgal J, Nayak PG, Pannakal ST, Priyadarsini KI, Unnikrishnan MK. Antidiabetic, antihyperlipidemic and antioxidant effects of the flavonoid rich fraction of Pilea microphylla (L.) in high fat diet/streptozotocin-induced diabetes in mice. Experimental and Toxicologic Pathology. 2012 Sep 1;64(6):651–8.

25. Bowe JE, Franklin ZJ, Hauge-Evans AC, King AJ, Persaud SJ, Jones PM. Metabolic phenotyping guidelines: assessing glucose homeostasis in rodent models. Journal of endocrinology. 2014 Sep 1;222(3):G13–25.

26. Kadam A, Mehta D, Jubin T, Mansuri MS, Begum R. Apoptosis inducing factor: Cellular protective function in Dictyostelium discoideum. Biochimica et Biophysica Acta (BBA)-Bioenergetics. 2020 Jan 25:148158.

27. Fontana L, Klein S. Aging, adiposity, and calorie restriction. Jama. 2007 Mar 7;297(9):986–94.

28. Huang P, Li S, Shao M, Qi Q, Zhao F, You J, Mao T, Li W, Yan Z, Liu Y. Calorie restriction and endurance exercise share potent anti-inflammatory function in adipose tissues in ameliorating diet-induced obesity and insulin resistance in mice. Nutrition & metabolism. 2010 Dec;7(1):1–9.

29. Baumeier C, Kaiser D, Heeren J, Scheja L, John C, Weise C, Eravci M, Lagerpusch M, Schulze G, Joost HG, Schwenk RW. Caloric restriction and intermittent fasting alter hepatic lipid droplet proteome and diacylglycerol species and prevent diabetes in NZO mice. Biochimica et Biophysica Acta (BBA)-Molecular and Cell Biology of Lipids. 2015 May 1;1851(5):566–76.

30. Smith JJ, Kenney RD, Gagne DJ, Frushour BP, Ladd W, Galonek HL, Israelian K, Song J, Razvadauskaite G, Lynch AV, Carney DP. Small molecule activators of SIRT1 replicate signaling pathways triggered by calorie restriction in vivo. BMC systems biology. 2009 Dec 1;3(1):31.

31. Rodgers JT, Lerin C, Haas W, Gygi SP, Spiegelman BM, Puigserver P. Nutrient control of glucose homeostasis through a complex of PGC-1α and SIRT1. Nature. 2005 Mar;434(7029):113–8.

32. Civitarese AE, Carling S, Heilbronn LK, Hulver MH, Ukropcova B, Deutsch WA, Smith SR, Ravussin E. Calorie restriction increases muscle mitochondrial biogenesis in healthy humans. PLoS med. 2007 Mar 6;4(3):e76.

33. Rathwa NN, Patel R, Pramanik S, Parmar NR, Ramachandran A, Begum R. Calorie Restriction in Combination with GABA Ameliorates Type 2 Diabetes. Diabetes 2019 Jun; 68 (Supplement 1): 143–LB

34. Oh SH, Moon YJ, Oh CH. γ-Aminobutyric acid (GABA) content of selected uncooked foods. Journal of Food Science and Nutrition. 2003 Mar;8(1):75–8.

35. Dhakal R, Bajpai VK, Baek KH. Production of GABA (γ-aminobutyric acid) by microorganisms: a review. Brazilian Journal of Microbiology. 2012 Dec;43(4):1230–41.

36. Wang S, Xiang YY, Zhu J, Yi F, Li J, Liu C, Lu WY. Protective roles of hepatic GABA signaling in acute liver injury of rats. American Journal of Physiology-Gastrointestinal and Liver Physiology. 2017 Mar 1;312(3):G208–18.

37. Ye DW, Rong XL, Xu AM, Guo J. Liver-adipose tissue crosstalk: A key player in the pathogenesis of glucolipid metabolic disease. Chinese journal of integrative medicine. 2017 Jun 1;23(6):410–4.

38. Al Nahdi AM, John A, Raza H. Elucidation of molecular mechanisms of streptozotocin-induced oxidative stress, apoptosis, and mitochondrial dysfunction in Rin-5F pancreatic β-cells. Oxidative Medicine and Cellular Longevity. 2017 Oct;2017.

39. Li K, Wu D, Chen X, Zhang T, Zhang L, Yi Y, Miao Z, Jin N, Bi X, Wang H, Xu J. Current and emerging biomarkers of cell death in human disease. BioMed research international. 2014 Jan 1;2014.

40. Hauke TJ. EXPRESSION PATTERN OF APOPTOSIS-INDUCING FACTOR IN THE KIDNEYS OF STREPTOZOTOCIN-INDUCED DIABETIC RATS (Doctoral dissertation, University of Split. School of Medicine. Department of Anatomy, Histology and Embryology).

