## Supplementary file for "Calorie Restriction in combination with GABA: improved glycaemic control in type 2 diabetes mouse model"

**Table S1.** Primers for the transcript levels.

| **Gene Primer** | **Sequence (5'-3')** | **Annealing**  **Temperature** | **Amplicon Size (bp)** | **Tissue** |
| --- | --- | --- | --- | --- |
| *Glucokinase* | **FP:**AGGAGGCCAGTGTAAAGATGT  **RP:**TCCCAGGTCTAAGGAGAGAAA | 56°C | 90bp | Liver |
| *PEPCK* | **FP:**CTGCATAACGGTCTGGACTTC  **RP:**CAGCAACTGCCCGTACTCC | 65°C | 151bp |  |
| *G6Pase* | **FP:**CTGTTTGGACAACGCCCGTAT  **RP:**AGGTGACAGGGAACTGCTTTA | 56°C | 91bp |  |
| *Glut2* | **FP:**CTTGGAAGGATCAAAGCAATG  **RP:**CAGTCCTGAAATTAGCCCAC | 60°C | 150bp |  |
| *Glycogen Synthase* | **FP:**ACCAAGGCCAAAACGACAG  **RP:**GGGCTCACATTGTTCTACTTG | 61°C | 102bp |  |
| *Glycogen Phosphorylase* | **FP:**GAGAAGCGACGGCAGATCA  **RP:**CTTGACCAGAGTGAAGTGCA | 65°C | 102bp |  |
| *SIRT-1* | **FP:**GATGAAGTTGACCTCCTCA  **RP:**GGGTATAGAACTTGGAATTAG | 64°C | 86bp | Liver, SK |
| *PGC-1α* | **FP:**AGCCGTGACCACTGACAACGA  **RP:**GTAGCTGAGCTGAGTGTTGGC | 69°C | 129bp |  |
| *TFAM* | **FP:**CTGAGGAAAAGCAGGCATA  **RP:**ATGTCTCCGGATCGTTTCAC | 69°C | 142bp |  |
| *ATGL* | **FP:**TGTGGCCTCATTCCTCCTAC  **RP:**TCGTGGATGTTGGTGGAGCT | 61.8°C | 158bp | AT |
| *ACC-1* | **FP:**ACGCTCAGGTCACCAAAAAGAAT  **RP:**GTAGGGTCCCGGCCACAT | 57°C | 70bp |  |
| *GAPDH* | **FP:** AGGTCGGTGTGAACGGATTTG  **RP:** TGTAGACCATGTAGTTGAGGT | 56°C | 123bp | Liver, SK, AT |

FP: Forward Primer; RP: Reverse Primer; bp: base pair; SK: Skeletal Muscle; AT: Adipose Tissue

**Table S2a: BW of control and HFD+STZ treated mice groups.**

| **B.W (g)** | **Control** | **HFD+STZ** | ***p*-value** |
| --- | --- | --- | --- |
| Week 1 | 22±1.4 | 23.9±1.4 | ns |
| Week 25 | 25±2.3 | 31.07±4.57 | ****p*<0.001 |

Data are presented as Mean ±SD. Statistical significance was considered at *p*<0.05. ns=non-significant

**Table S2b: FBG of control and HFD+STZ treated groups.**

| **FBG (mg/dL)** | **Control** | **HFD+STZ** | ***p*-value** |
| --- | --- | --- | --- |
| Week 1 | 139±12 | 138±22.17 | ns |
| Week 25 | 140±57.7 | 362.5±45.6 | ****p*<0.001 |

Data are presented as Mean ±SD. Statistical significance was considered at *p*<0.05. ns=non-significant

**Table S3a. The transcript levels of gluco-regulatory genes in liver.**

| **Gene** | **Treatment** | **Fold Change** | ***p*-value** |
| --- | --- | --- | --- |
| ***Glucokinase*** | HFD+STZ | 0.04 | - |
|  | CR | 0.41 | ns |
|  | GABA | 0.16 | ns |
|  | CR+GABA | 0.5 | ** |
| ***PEPCK*** | HFD+STZ | 5.61 | - |
|  | CR | 3.4 | * |
|  | GABA | 1.04 | ** |
|  | CR+GABA | 2.77 | ** |
| ***G6Pase*** | HFD+STZ | 2.32 | - |
|  | CR | 0.24 | * |
|  | GABA | 0.18 | * |
|  | CR+GABA | 0.28 | * |
| ***Glut2*** | HFD+STZ | 0.75 | - |
|  | CR | 0.57 | ** |
|  | GABA | 0.32 | ** |
|  | CR+GABA | 0.36 | ** |
| ***Glycogen Synthase*** | HFD+STZ | 0.01 | - |
|  | CR | 0.13 | ns |
|  | GABA | 0.28 | ns |
|  | CR+GABA | 0.48 | ns |
| ***Glycogen***  ***Phosphorylase*** | HFD+STZ | 2.31 | - |
|  | CR | 0.43 | * |
|  | GABA | 0.08 | * |
|  | CR+GABA | 0.17 | * |
| ***SIRT-1*** | HFD+STZ | 1.4 | - |
|  | CR | 3.2 | ns |
|  | GABA | 1.6 | ns |
|  | CR+GABA | 6.0 | ** |
| ***PGC-1α*** | HFD+STZ | 1.1 | - |
|  | CR | 4.4 | ** |
|  | GABA | 2.4 | ns |
|  | CR+GABA | 9.2 | *** |

Data are presented as Mean ±SD. (ns=non-significant, **p*<0.05, ***p*<0.01,

****p*<0.001, n=5/ group).

**Table S3b. The transcript levels of lipid metabolism marker genes in adipose tissue.**

| **Gene** | **Treatment** | **Fold Change** | ***p*-value** |
| --- | --- | --- | --- |
| ***ATGL*** | HFD+STZ | 2.96 | - |
|  | CR | 3.4 | ns |
|  | GABA | 4.4 | ns |
|  | CR+GABA | 8.04 | * |
| ***ACC-1*** | HFD+STZ | 3.55 | - |
|  | CR | 15.37 | ns |
|  | GABA | 10.11 | ns |
|  | CR+GABA | 16.25 | * |
| ***FOXO*** | HFD+STZ | 1.3 | - |
|  | CR | 1.8 | ns |
|  | GABA | 2.6 | ns |
|  | CR+GABA | 5.1 | * |

Data are presented as Mean ±SD. (ns=non-significant, **p*<0.05,

***p*<0.01, ****p*<0.001, n=5/ group).

**Table S3c. The transcript levels of mitochondrial biogenesis genes in skeletal muscle.**

| **Gene**  **(In skeletal muscle)** | **Treatment** | **Fold Change** | ***p*-value** |
| --- | --- | --- | --- |
| ***SIRT-1*** | HFD+STZ | 0.16 | - |
|  | CR | 3.27 | ns |
|  | GABA | 2.34 | ns |
|  | CR+GABA | 5.7 | *** |
| ***PGC-1α*** | HFD+STZ | 0.17 | - |
|  | CR | 4.73 | * |
|  | GABA | 0.9 | ns |
|  | CR+GABA | 5.8 | *** |
| ***tFAM*** | HFD+STZ | 1.85 | - |
|  | CR | 2.63 | ns |
|  | GABA | 0.84 | ns |
|  | CR+GABA | 3.79 | *** |

Data are presented as Mean ±SD. (ns=non-significant, **p*<0.05,

***p*<0.01, ****p*<0.001, n=5/ group).

**Table S4. Antibodies for the IHC studies.**

| **Primary Antibody** | **Secondary Antibody** | **Excitation (nm)** | **Emission (nm)** |
| --- | --- | --- | --- |
| Anti-Insulin  (1:100, Guinea Pig)  [DAKO Agilent, USA] | Alexa 488 (1:500, Donkey)  [Jackson ImmunoResearch Laboratories, Inc. USA] | 493 | 519 |
| NGN3 (1:50, rabbit)  [Thermo Fisher Scientific, USA] | Alexa 647 (1:500, Donkey)  [Jackson ImmunoResearch Laboratories, Inc. USA] | 651 | 667 |
| ARX (1:500, Rabbit)  [Sigma-Aldrich, Germany] |  |  |  |
| Anti-Glucagon (1:200, rabbit) [Cell Signaling Technology, USA] |  |  |  |
| Anti-AIF (1:400, rabbit) [Thermo Fisher Scientific, USA] |  |  |  |
| PDX-1 (1:1000, Goat) [Abcam, USA] | Rhodamine Red (1:200, Donkey)  [Jackson ImmunoResearch Laboratories, Inc. USA] | 570 | 590 |
| PAX-4 (1:500, Goat)  [Sigma-Aldrich, Germany] |  |  |  |
| BrdU (1:100, Goat)  [Abcam, USA] | Rhodamine Red (1:200, Rat)  [Jackson ImmunoResearch Laboratories, Inc. USA] | 570 | 590 |

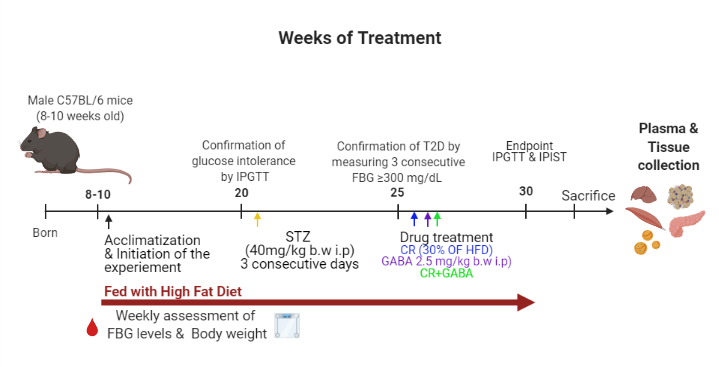

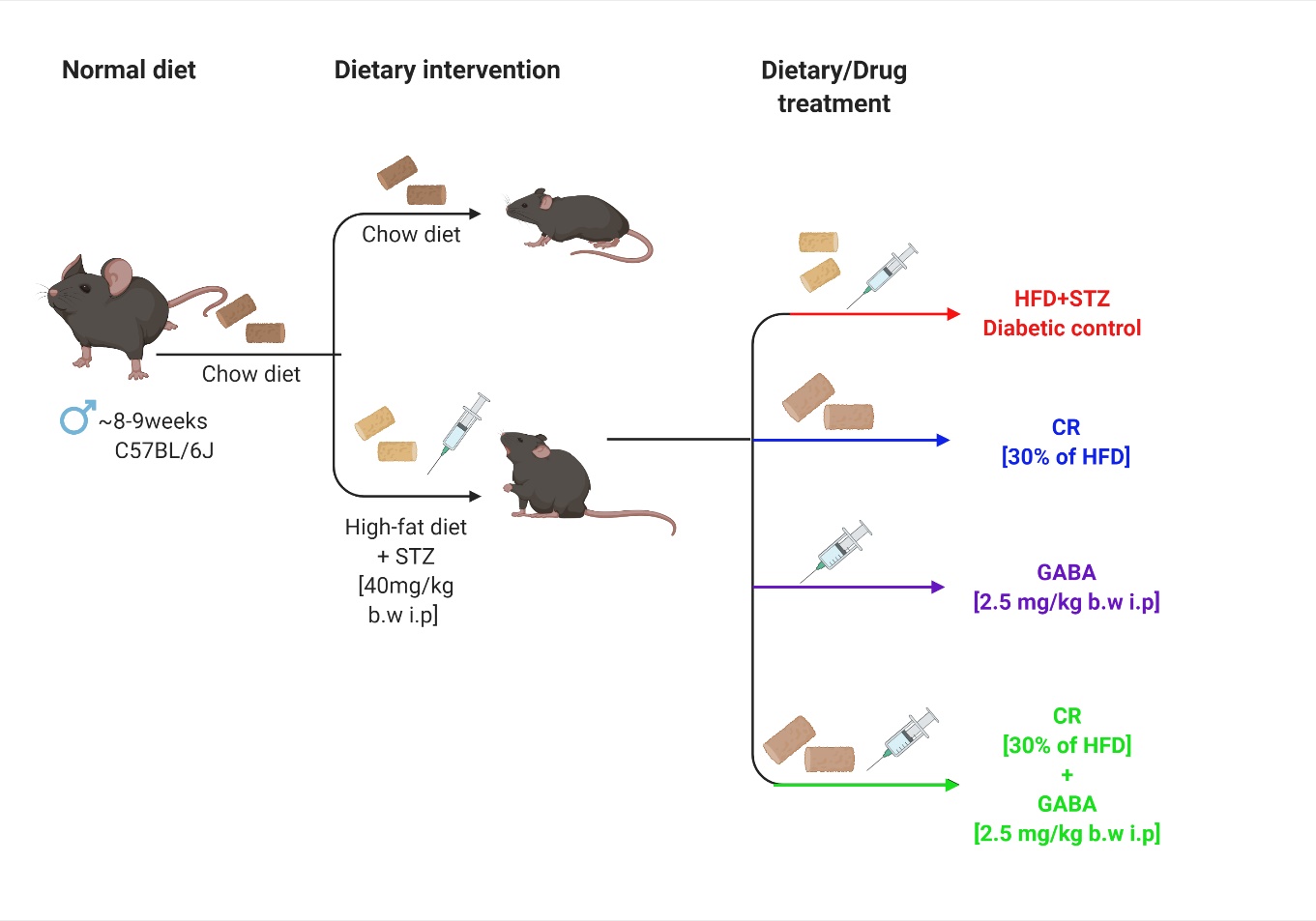
**Figure S1. The experimental timeline:** ~8-9 weeks old male C57BL/6 mice were procured and subjected to 1 week acclimatization. 25 of them were fed with HFD for 20 weeks to induce obesity and insulin resistance while 6 were fed with chow diet. Food intake, water consumption, BW and FBG levels were monitored weekly. After 20 weeks, the HFD fed mice received three low consecutive doses of STZ (40 mg/kg i.p.) to induce β-cells loss. Once obesity-induced T2D was confirmed, these animals were divided randomly into four groups for a treatment of 6 weeks. Post treatment, IPGTT and IPIST was carried out. The animals were sacrificed and tissues were harvested for further analysis.

**Figure S2. Body weight and fasting blood glucose levels in experimental diabetic mice** (Control and post- HFD+STZ treatment). A significant difference in BW and FBG was observed in the animals before and after HFD+STZ treatment. (**p<0.01, ***p<0.001 respectively) (Control, n=5 and post HFD+STZ, n=25)

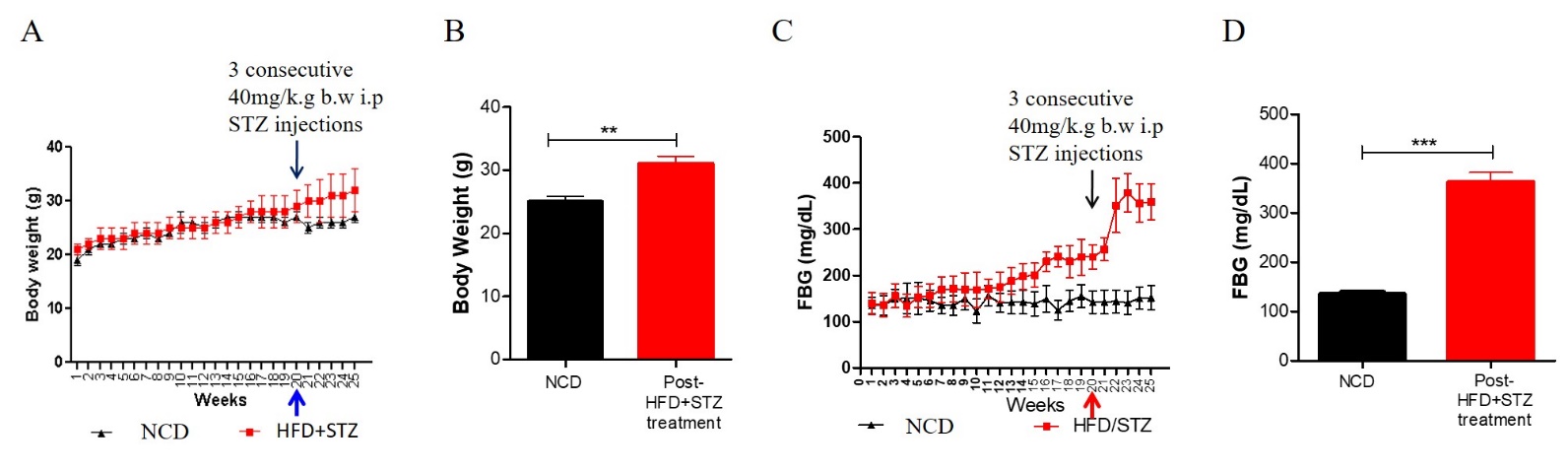
